# LGR4 is essential for maintaining β-cell homeostasis through suppression of RANK

**DOI:** 10.1101/2024.05.10.593645

**Authors:** Joanna Filipowska, Zelda Cisneros, Nancy Leon-Rivera, Peng Wang, Randy Kang, Geming Lu, Yate-Ching Yuan, Supriyo Bhattacharya, Sangeeta Dhawan, Adolfo Garcia-Ocaña, Nagesha Guthalu Kondegowda, Rupangi C. Vasavada

**Affiliations:** Arthur Riggs Diabetes and Metabolism Research Institute, City of Hope, Duarte, CA 91010, USA; Department of Translational Research and Cellular Therapeutics, City of Hope, Duarte, CA 91010, USA; Diabetes, Obesity and Metabolism Institute, and Division of Endocrinology, Diabetes and Bone Diseases, Icahn School of Medicine at Mount Sinai, New York, NY 10029, USA; Department of Molecular and Cellular Endocrinology, City of Hope, Duarte, CA 91010, USA; Division of Research Informatics, City of Hope, Duarte, CA 91010, USA; Department of Molecular Imaging and Therapy, City of Hope, Duarte, CA 91010, USA

## Abstract

Pancreatic β-cell stress contributes to diabetes progression. This study demonstrates that Leucine-rich repeat-containing G-protein-coupled-receptor-4 (LGR4) is critical for maintaining β-cell health and is modulated by stressors. *In vitro*, *Lgr4* knockdown decreases proliferation and survival in rodent β-cells, while overexpression protects against cytokine-induced cell death in rodent and human β-cells. Mechanistically, LGR4 suppresses Receptor Activator of Nuclear Factor Kappa B (NFκB) (RANK) and its subsequent activation of NFκB to protect β-cells. β-cell-specific *Lgr4*-conditional knockout (cko) mice exhibit normal glucose homeostasis but increased β-cell death in both sexes and decreased proliferation only in females. Male *Lgr4*cko mice under stress display reduced β-cell proliferation and a further increase in β-cell death. Upon aging, both male and female *Lgr4*cko mice display impaired β-cell homeostasis, however, only female mice are glucose intolerant with decreased plasma insulin. We show that LGR4 is required for maintaining β-cell health under basal and stress-induced conditions, through suppression of RANK.

**Teaser:** LGR4 receptor is critical for maintaining β-cell health under basal and stressed conditions, through suppression of RANK.

## Introduction

Loss of functional insulin-producing β-cells and the inability to regenerate them play a key role in both Type 1 (T1D) and Type 2 (T2D) diabetes pathogenesis (1). Although T1D is an immune cell-mediated disease, recent literature indicates that β-cell stress could be a trigger for the autoimmune response (2, 3). Islet inflammation and β-cell stress are also hallmarks of T2D (4). Thus, identifying the regulators that maintain β-cell health and understanding their function is crucial in treating both types of diabetes. G-protein-coupled receptors (GPCRs) are therapeutic targets for multiple diseases as they comprise the largest group of receptors regulating key cellular processes in all tissues (5, 6). The glucagon-like peptide 1 receptor (GLP1R) is a prime example of a GPCR targeted for therapeutic treatment of diabetes (7, 8).

Leucine-rich repeat-containing G protein-coupled receptor 4 (LGR4), a member of the GPCR subfamily B, is the fourth most abundant GPCR in human islets, with a predominant expression in β compared to α cells (9). LGR4 is expressed in rodent and human pancreatic β-cells (9, 10). Classically, LGR4 binds R-spondins (RSPO) to potentiate Wnt signaling, particularly, during embryonic development (patterning and stem cell differentiation) (11–14). Recently, another ligand, Receptor Activator of Nuclear Factor Kappa B (NFκB) (RANK) ligand (RANKL), was identified to bind LGR4 receptor in osteoclast precursor cells (15). In osteoclasts, RANKL binding to its classical receptor, RANK induces osteoclast activation and bone loss (16–18). However, the same ligand, RANKL, binding to the LGR4 receptor has the opposite effect, that of bone formation, by antagonizing RANKL/RANK activation (15). Our prior work has established that activation of the RANKL/RANK pathway inhibits β-cell proliferation and promotes cytokine-induced β-cell death and dysfunction (19, 20). Although LGR4 is expressed in β-cells (9, 10), the role of this receptor and its significance as a potential antagonist of RANK remains unknown. We hypothesized that LGR4 is required to suppress the negative effects of RANK on β-cell health, and therefore, reduced LGR4 levels would be detrimental to β-cell health.

LGR4 exhibits regenerative, anti-inflammatory, and anti-apoptotic functions across many tissues, including heart, skin, liver, kidney, and bone (14, 21–25). However, the functional significance of this receptor in β-cells remains unknown. We demonstrate that *Lgr4* levels are reduced in response to a variety of physiological and diabetogenic stressors, in rodent and human islets, as well as β-cells. Using *in vitro* loss and gain of function studies, we establish that LGR4 exerts a protective effect on β-cell health. Mechanistically, we reveal that LGR4 enhances β-cell survival through inhibiting RANK-Tumor necrosis factor receptor associated factor 6 (TRAF6) interaction and subsequent downstream NFκB activation. Through the generation of β-cell-specific *Lgr4*-conditional knockout (cko) mice we demonstrate that LGR4 is essential for maintaining normal β-cell replication and survival under basal and stressed conditions *in vivo*. Taken together, our studies identify LGR4 as a critical GPCR for preserving β-cell health, which acts by antagonizing the RANK pathway, known to be detrimental for β-cells, highlighting possible therapeutic implications of LGR4 in the context of diabetes-related β cell stress.

## Results

### LGR4 mRNA levels decrease in mouse and human islets and human β-cells under stress

Our analysis of published single-cell mouse islet transcriptome data (26) showed that *Lgr4* is expressed in the four major endocrine cell types of mouse pancreatic islets (Fig. 1A). We next examined *Lgr4* expression in mouse islets, in the context of various T2D and T1D relevant stressors. Meta-analysis of RNA-seq data of islets from young adult mice subjected to a high fat diet (HFD) for 2-months (27) (Fig. 1B) or from the leptin receptor-deficient db/db mice (28) (Fig. 1C) showed decreased expression of *Lgr4* compared to their respective controls, in a manner similar to multiple β-cell maturity markers. qPCR analysis of islets from 4- and 24-month-old aged wild type C57BL/6J male mice showed a similar reduction in *Lgr4* mRNA levels with aging (Fig. 1D). We also observed a significant decrease in *Lgr4* mRNA levels upon treatment of wild type mouse islets with proinflammatory cytokines for 24h (Fig. 1E), as assessed by qPCR. Analysis of published human islet single-cell transcriptome data (29) showed *LGR4* expression in all four major endocrine cell-types, with greater abundance in β-cells (Fig. 1F). As with mouse islets, proinflammatory cytokine-treatment of human islets for 24h led to a decrease in *LGR4* mRNA, based on our bulk RNA-seq analysis, along with a significant increase in inflammation markers (Fig. 1G, H) and this was confirmed independently on a separate cohort of human islets by qPCR analysis (Fig. 1I). To determine if cytokines decreased *LGR4* expression specifically in β-cells, we analyzed the published transcriptome of control (vehicle-treated) and cytokine-treated human EndoC-βH5 β-cell line (30) and found a significant decrease in *LGR4* mRNA levels 48h post cytokine-treatment (Fig. 1J).

**Figure 1.**
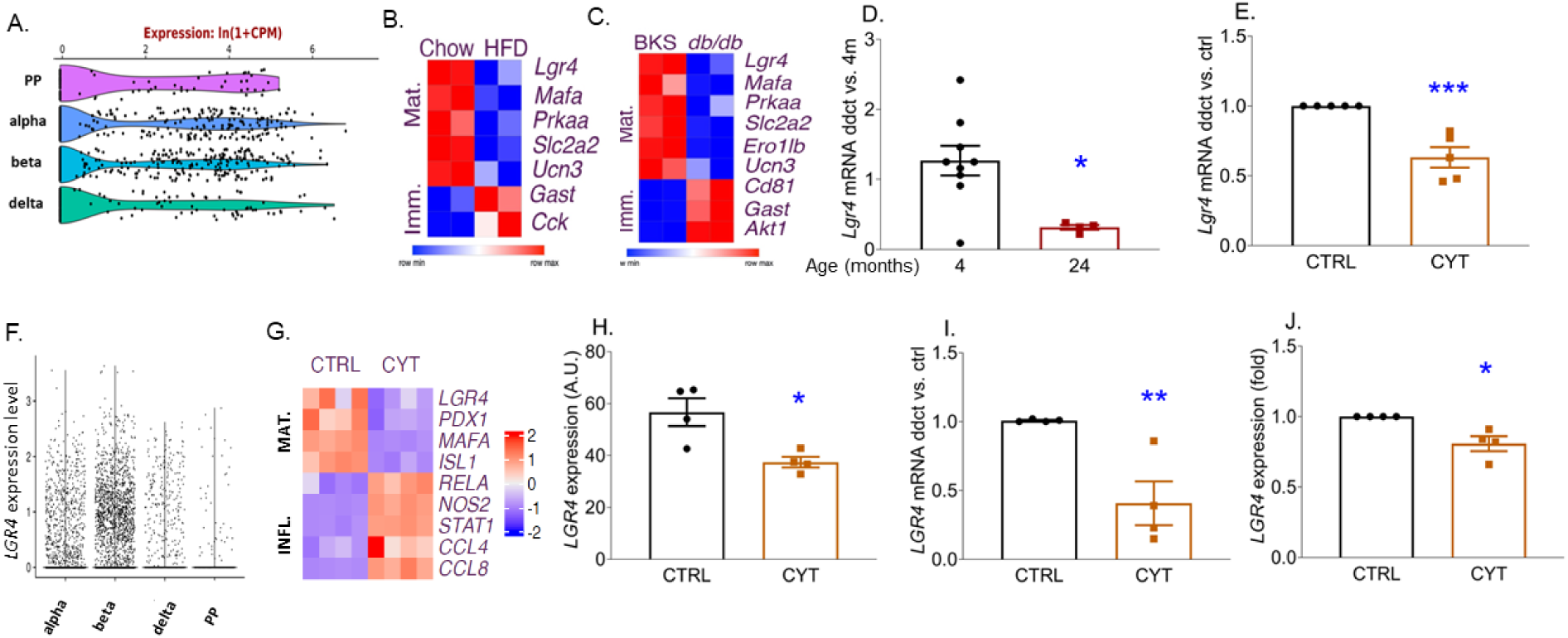
Lgr4 expression and regulation in mouse and human islets under stress. **(A)** Violin plot representation of *Lgr4* expression in the endocrine cells of mouse pancreatic islets based on published single-cell transcriptome data (26) analyzed using https://tabula-muris.ds.czbiohub.org/, with the different cell type clusters labeled. **(B, C)** Heat maps showing *Lgr4* mRNA, along with other maturity (mat) and immaturity (imm) markers in a meta-analysis of published RNA-seq data in islets from (B) adult (10-week-old) C57BL/6J mice fed with a chow diet or a high fat diet (HFD) for two months (n=2) (27) and (C) 8 weeks old db/db mice and age-matched BKS controls (n=2) (28). Blue and red colors indicate relative, low and high expression levels, respectively. qPCR analysis of *Lgr4* expression relative to *Cyclophilin A* as housekeeping gene in **(D)** mouse islets from 4- and 24-month-old C57Bl/6 male mice (n=4-9), **(E)** mouse islets treated with vehicle (CTRL) or cytokine mix (CYT) for 24h (n=5). **(F)** Violin plot representation of *LGR4* expression in the endocrine cells of human pancreatic islets based on published single-cell transcriptome analysis (29), with the different endocrine cell type clusters labeled. **(G)** Heat map showing *LGR4* mRNA, along with islet maturity (MAT) and inflammatory (INFL) markers in RNA- seq analysis of human islets treated with vehicle (CTRL) or cytokine mix (CYT) for 24h (n=4). Blue and red colors indicate relative, low and high expression, respectively. **(H)** *LGR4* mRNA levels in arbitrary units (A.U.) from the RNA-seq analysis of human islets described in (G). **(I)** qPCR analysis of *LGR4* expression relative to *BETA ACTIN* in an independent cohort of human islets treated with vehicle (CTRL) or cytokine mix (CYT) for 24h (n=4). **(J)** *LGR4* expression in the human β-cell line (EndoC-βH5) treated with vehicle (CTRL) or CYT for 48h based on meta-analysis of published transcriptome data (30) using https://www.ncbi.nlm.nih.gov/geo/query/acc.cgi?acc=GSE218735 (n=4). *p<0.05, **p<0.01, ***p<0.001 vs 4 month or CTRL. Individual symbols in the graphs represent individual mouse islet preps (D, E), or independent experiments on different human islet preps (H-J). All data represent mean ± SEM. Statistical analysis was done by t-test.

### *Lgr4* downregulation leads to decreased proliferation and increased β-cell death in basal and stress-induced conditions in INS1 cells and mouse islet cells *in vitro*

To discern the role of LGR4 in β-cells, we used *Lgr4*-siRNA in INS1 cells to induce an ∼50% reduction in *Lgr4* mRNA compared to control (untreated cells) or scrambled (SC)-siRNA transfected cells (Fig. 2A). INS1 cell proliferation, as assessed by bromodeoxyuridine (BrdU) incorporation (Fig. 2B), was significantly decreased in basal conditions, in *Lgr4* knockdown (38.42±3.44%), relative to untreated control (48.28±2.41%) and SC-siRNA (49.83±1.40%)-treated cells (Fig. 2C). On the other hand, cell death measured by Cleaved Caspase-3 staining, was significantly increased more than three-fold in *Lgr4*-deficient versus untreated or SC-siRNA treated INS1 cells under basal conditions (Fig. 2D). Next, we tested if *Lgr4* depletion exacerbates cytokine-induced cell death. Indeed, the increased cell death observed upon proinflammatory cytokine-treatment in untreated control and SC-siRNA treated INS1 cells was further significantly increased at 16h and 24h in *Lgr4*-deficient cells (Fig. 2E). To examine the effect of LGR4 in primary rodent β-cells *in vitro*, we transduced mouse islet cells from *Lgr4*-floxed mice with Adenovirus-Cre recombinase (Ad-*Cre*) to knockdown *Lgr4* or Ad-*LacZ* as a control. Co-staining for insulin and Cre showed robust Cre expression in the Ad-*Cre* transduced β-cells compared to Ad-*LacZ* control transduction (Fig. 2F). We observed a 50% reduction in *Lgr4* mRNA, 72h after transduction, in the Ad-*Cre*-versus Ad-*LacZ*-transduced mouse islet cells from both male and female mice (Fig. 2G). Interestingly, acute *Lgr4* knockout (ko) impacted β-cell proliferation, as assessed by phospho-histone 3 (pHH3)-insulin co-staining (Fig. 2H), in a sex-specific manner. While *Lgr4*ko significantly decreased β-cell replication by ∼50% in islets from female mice (Fig. 2I) compared to controls, it did not impact β-cell replication in islets from male mice (Fig. 2J). On the other hand, β-cell death, assessed by TUNEL-insulin co-staining significantly increased in both male and female mouse islets upon acute *Lgr4*ko *in vitro* (3.51±0.21%) compared to controls (1.93±0.21%) (Fig. 2K).

**Figure 2.**
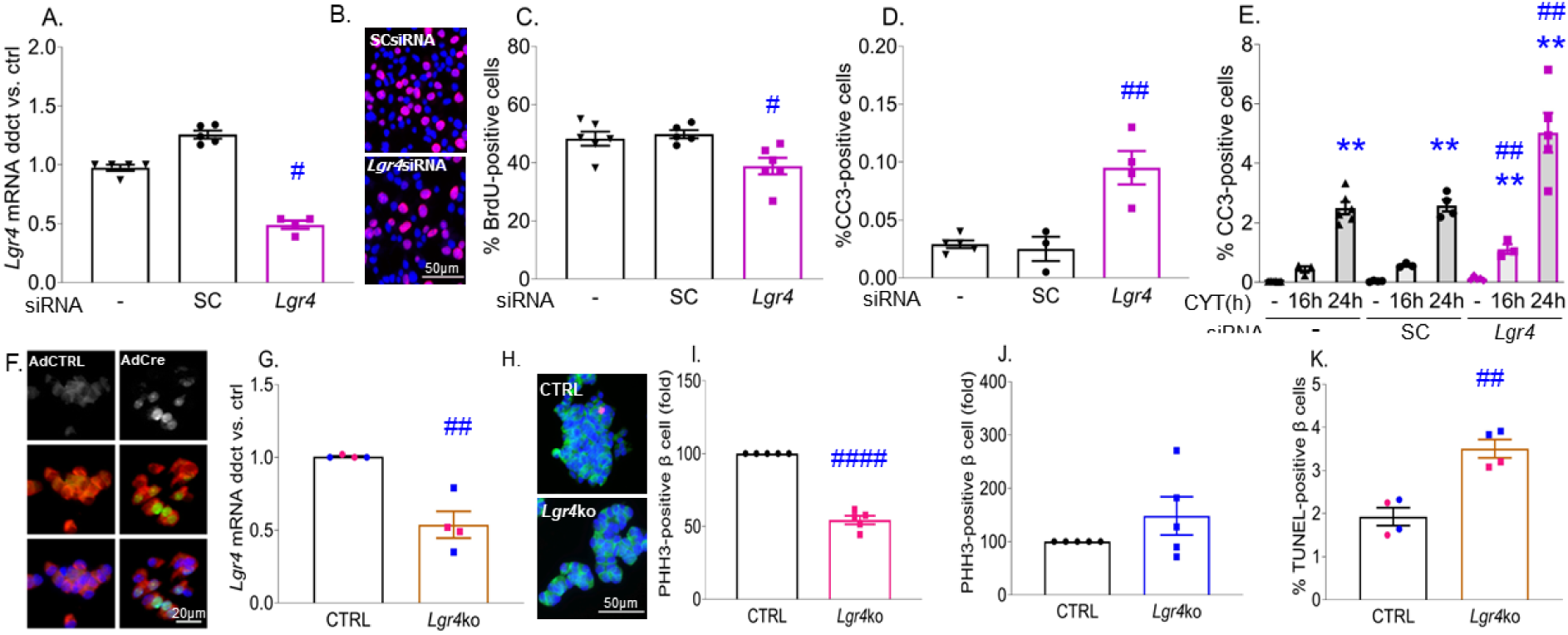
*Lgr4* downregulation decreases β-cell proliferation and increases β-cell death in INS-1 cells and mouse islets *in vitro*. INS-1 cells un-transfected (-) or transfected with scrambled (SC) or *Lgr4* siRNA for 48-72h were assayed for **(A)** *Lgr4* expression relative to Cyclophilin A by qPCR after 48h (n=4-5); **(B)** BrdU incorporation by immunofluorescent staining for BrdU (red) and nuclear DAPI (blue), after 72h, with the addition of BrdU to the culture medium for the last 2h, scale bar is 50µm; **(C)** proliferation measured as percent BrdU-positive cells for the treatment described in (B) (n=6); **(D, E)** cell death measured as percent Cleaved Caspase-3 (CC3)-positive cells (D) basally after 72h (n=3-5), or (E) after treatment without (-) or with cytokine mix (CYT) for the last 16 or 24h (n=3-6). Mouse islet cells transduced with adenovirus (Ad) *LacZ* as control (CTRL) or Ad*Cre* (*Lgr4*ko) for 72h and assayed for **(F)** Cre recombinase (green or white), insulin (red) and DAPI (blue) by immunofluorescent staining after 72h shown as separate or merged panels, scale bar is 20µm; **(G)** *Lgr4* relative to *Cyclophilin A* mRNA by qPCR after 72h, with blue symbols representing male and pink symbols representing female mouse islets (n=4); **(H, I, J)** β-cell proliferation after 72h, on (H) pHH3 (red), insulin (green) and DAPI (blue) stained immunofluorescent cells, scale bar is 50µm, measured as percent pHH3 and insulin double-positive cells and depicted as fold over CTRL in (I) male (n=5), (CTRL 0.28±0.08% proliferating β cells) and (J) female (n=5) mice, (CTRL 0.89±0.23% proliferating β cells); **(K)** percent β-cell death measured as TUNEL- and insulin-positive cells, after 72h, with blue symbols representing male and pink symbols representing female mouse islets (n=4). ^#^p<0.05, ^##^p<0.01, ^####^p<0.0001 vs un-transfected (-) and SC groups for the same treatment in INS-1 cells, and vs CTRL for mouse islets. **p<0.01 vs untreated (-) INS-1 cells from the same group. Individual symbols in the graphs represent independent experiments in INS-1 cells (A, C-E) or individual mouse islet preps (G, I-K), averaging duplicate samples for all experiments. All data represent mean ± SEM. Statistical analysis was by t-test for comparison of two groups, and by ANOVA with Tukey’s post-hoc analysis for comparison of more than two groups.

### LGR4 overexpression protects β-cells against proinflammatory cytokine-induced cell death in INS1 cells, mouse, and human islets

Based on the effects of *Lgr4* deficiency, we sought to determine if *Lgr4* overexpression would exert a protective effect on β-cells. *Lgr4* expression was increased two-fold in Ad-*Lgr4*-transduced when compared to un-transduced or Ad-Ctrl-transduced INS1 cells (Fig. 3A). Ad-*Lgr4* transduction significantly reduced proinflammatory cytokine-induced cell death (2.94±0.35%), when compared to un-transduced (4.89±0.73%) or Ad-Ctrl (5.79±0.57%) transduced cells, as assessed by Cleaved Caspase-3 staining (Fig. 3B, C). Next, we used mouse islet cells transduced with species-specific Ad-*Lgr4* to induce *Lgr4* expression (Fig. 3D). Ad-*Lgr4* was able to reduce cytokine-induced β-cell death by 50% relative to Ad-Ctrl transduced cells, as measured by TUNEL-insulin co-staining (Fig. 3E, F). Notably, human islet cells transduced with species-specific Ad-*LGR4* demonstrated a significant increase in *LGR4* mRNA relative to the Ad-Ctrl-transduced cells (Fig. 3G), corresponding to an increase in LGR4 protein levels as assessed by insulin-LGR4 co-staining (Fig. 3H). Human β-cells were protected against proinflammatory cytokine-induced cell death under *LGR4* overexpression compared to Ad-Ctrl transduced cells (1.61±0.42-versus 4.86±1.36-fold increase in cell death compared to vehicle-treated Ad-Ctrl cells, respectively) (Fig. 3I). However, LGR4 overexpression did not affect the low level of basal cell death observed in INS1, mouse or human β-cells (Fig. 3C, F, I).

**Figure 3.**
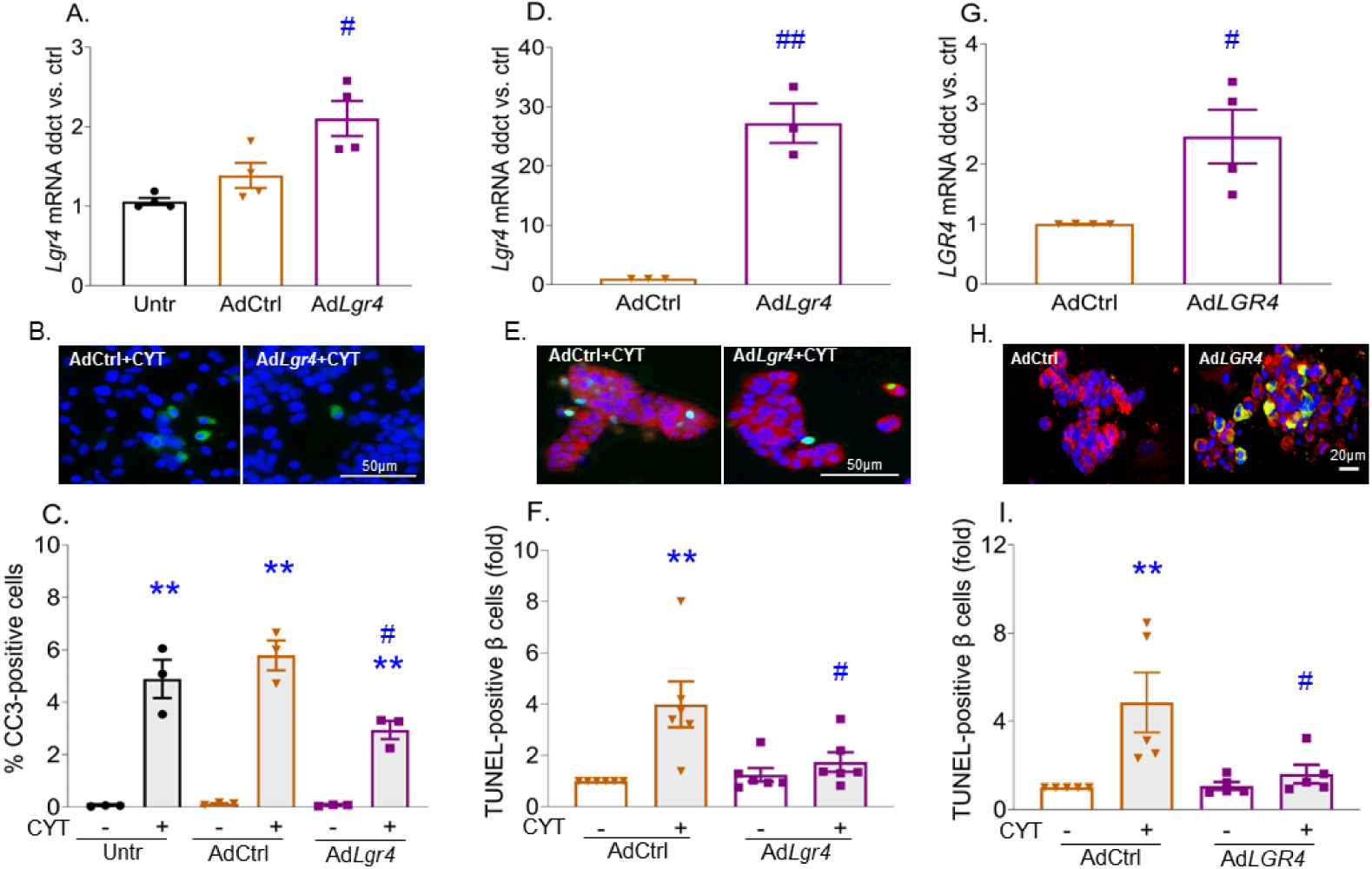
LGR4 overexpression protects β-cells against proinflammatory cytokine-induced cell death in INS1 cells, mouse, and human islets. **(A-C)** INS-1 cells were untreated (Untr) or transduced with adenovirus (Ad) *LacZ* as control (AdCtrl) or Ad*Lgr4* and (A) assessed for *Lgr4* mRNA expression by qPCR after 48h (n=4); or treated after 24h without (-) or with cytokine mix (CYT) for an additional 24h and (B) stained for Cleaved Caspase-3 (CC3) (green) and DAPI (blue), and (C) quantified for cell death as percent CC3-positive cells (n=3). **(D-F)** Mouse or **(G-I)** human islet cells were transduced with adenovirus (Ad) *Cre* as control (AdCtrl) or species-specific Ad*Lgr4* and (D, G) assessed for *Lgr4* mRNA expression by qPCR after 48h in (D) mouse (n=3) and (G) human (n=4) islets; or treated after 24h without (-) or with cytokine mix (CYT) for an additional 24h. Mouse islet cells were (E) stained for TUNEL (green), insulin (red), and DAPI (blue), and (F) quantified for percent β-cell death represented as fold over Ad-Ctrl (n=6), (Ad-Ctrl/- 1.30±0.56% TUNEL-positive β cells). Human islet cells were (H) stained for LGR4 (green), insulin (red) and DAPI (blue) after 72h; and (I) quantified for percent β-cell death represented as fold over Ad-Ctrl (n=5), (Ad-Ctrl/- 0.68±0.21% TUNEL-positive β cells). ^#^p<0.05, ^##^p<0.01 vs Untr and AdCtrl groups in INS-1 cells, and vs AdCtrl group in mouse and human islets with similar treatments. **p<0.01 vs no cytokine treatment (-) from the same group. White bar indicates the scale for the immunofluorescent images (50µm). Individual symbols in the graphs represent independent experiments in INS1 cells (A, C) or individual mouse (D, F) or human (G, I) islet preps, averaging duplicate samples for all experiments. All data represent mean ± SEM. Statistical analysis was by t-test for comparison of two groups, and by ANOVA with Tukey’s post-hoc analysis for comparison of more than two groups.

### LGR4 improves β-cell survival through the suppression of RANK-TRAF6 interaction and subsequent downstream activation of NFκB

We have previously demonstrated that cytokine-induced β-cell death requires RANK-TRAF6 interaction which leads to NFκB pathway activation (20). In osteoclasts, LGR4 binds RANKL thereby blocking the induction of the RANKL/RANK pathway, and thus reducing osteoclast activation (15). We hypothesized that LGR4 mediates its protective effects on β-cell survival through inhibition of the RANK pathway. To test this, we performed a rescue experiment where both *Lgr4* and *Rank* were knocked down simultaneously in INS1 cells, to examine the effect of the double ko on basal cell death. Indeed, *Lgr4*-siRNA and *Rank*-siRNA transfected singly or simultaneously in INS1 cells induced a significant (∼50%) reduction in *Lgr4* (Fig. 4A) and *Rank* (Fig. 4B) mRNAs, respectively, when compared to control SC-siRNA transfected cells. As expected, basal INS1 cell death was significantly increased with *Lgr4*-siRNA alone (0.20±0.05%) compared to SC-siRNA (0.06±0.01%), with no effect of *Rank*-siRNA alone (0.04±0.01%). However, double ko of *Lgr4* and *Rank* significantly reduced β-cell death (0.05±0.00%) to control levels observed in SC-siRNA-transfected cells (Fig. 4C). Thus, simultaneous knockdown of *Rank* and *Lgr4* rescued the increased cell death phenotype observed with Lgr4 knockdown alone, implying that LGR4 mediates its beneficial effects on β-cell survival through inhibition of RANK under basal conditions.

**Figure 4.**
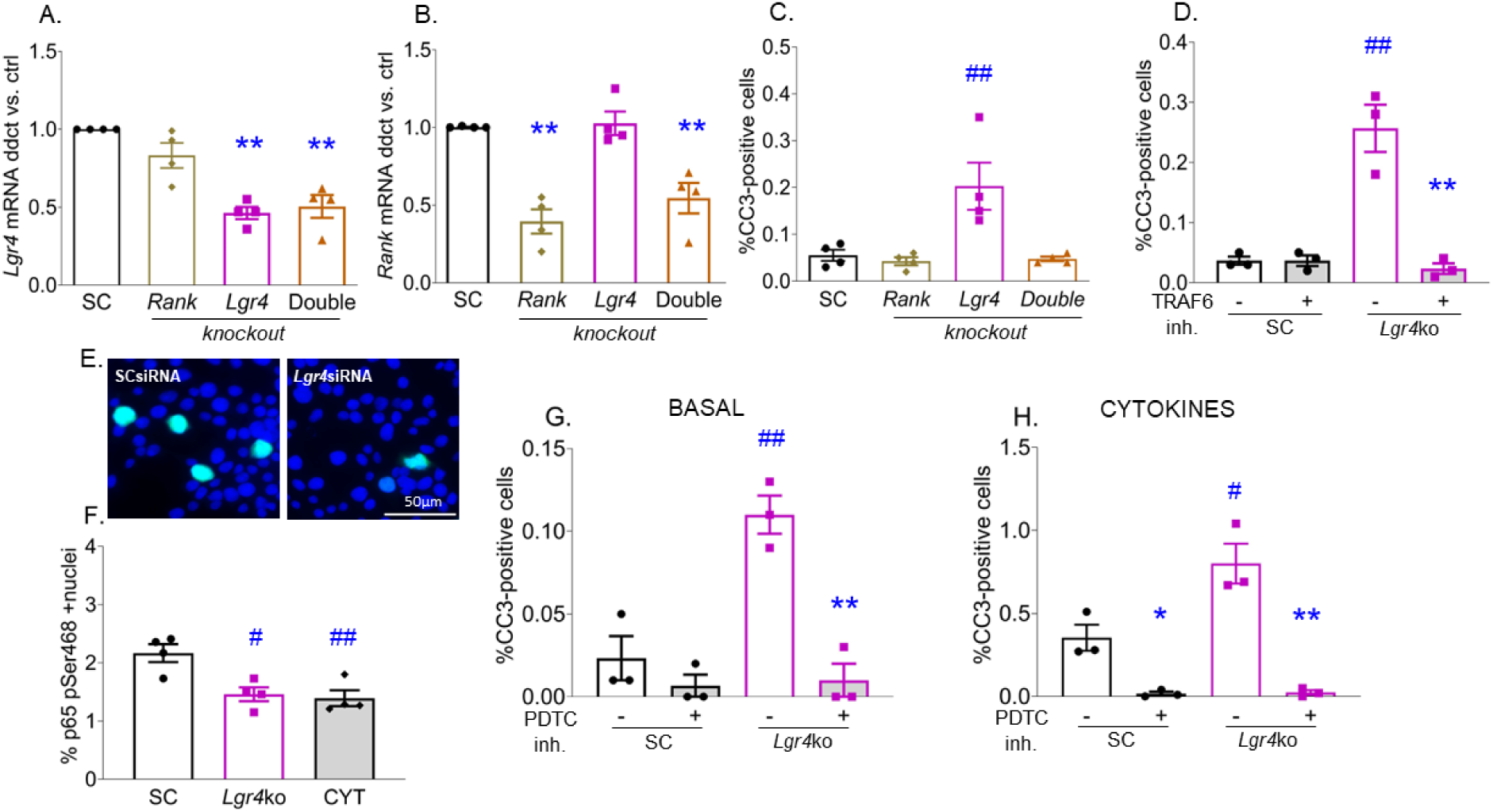
LGR4 improves β-cell survival through suppressing RANK-TRAF6 interaction and the subsequent activation of NFκB. INS-1 cells were transfected with scrambled (SC), *Lgr4* (*Lgr4*ko) or *Lgr4* and *Rank* (Double ko) siRNAs and assayed for **(A)** *Lgr4* and **(B)** *Rank* mRNAs by qPCR after 48h; **(C)** cell death by percent CC3-positive cells after 48h (n=4); **(D)** percent CC3-positive cells after treatment with (+) or without (-) TRAF6 peptide inhibitor for the last 16h (n=4); **(E)** phSer468 NFκB p65 (green) and DAPI (blue) by immunofluorescent staining; **(F)** nuclear phSer468 NFκB quantification, with cytokine (CYT) treatment used as a positive control (n=4); **(G-H)** percent CC3-positive cells without (-) or with (+) treatment with PDTC inhibitor (G) under basal (n=3) and (H) after cytokine-treatment for 16h (n=3). ^#^p<0.05, ^##^p<0.01 vs SC and Double ko groups with the same treatment. *p<0.05, **p<0.01 vs no inhibitor treatment (-) from the same group. White bar indicates the scale for the immunofluorescent images (50µm). Individual symbols in the graphs represent independent experiments in INS1 cells, averaging duplicate samples. All data represent mean ± SEM. Statistical analysis was by t-test for comparison of two groups, and by ANOVA with Tukey’s post-hoc analysis for comparison of more than two groups.

We previously showed that RANK/TRAF6 interaction is critical for inducing NFκB activation and cytokine-induced β-cell death (20). To determine if the RANK/TRAF6 interaction mediates the increased cell death in Lgr4-deficient INS1 cells, we treated these cells with a peptide inhibitor that specifically blocks RANK-TRAF6 interaction (31). Indeed, the increased cell death observed in *Lgr4*-deficient INS1 cells was significantly reduced to control levels with RANK-TRAF6-peptide inhibitor treatment (Fig. 4D), underscoring the importance of RANK-TRAF6 interaction in mediating the increased cell death observed in *Lgr4* knockdown conditions. To assess the role of NFκB, we stained for NFκB p65 (RELA) phosphorylation at Serine-468 (Fig. 4E), a marker for NFκB pathway deactivation, as it allows degradation of NFκB (32–34). Indeed, *Lgr4*-deficient INS1 cells exhibited a significant decrease in NFκB-phSer468 in basal conditions compared to SC-siRNA transfected cells (Fig. 4F), implying activation of the NFκB pathway upon *Lgr4* knockdown. The decrease in NFκB-phSer468 with *Lgr4*-deficiency was similar to that observed with cytokines (Fig. 4F), used as a positive control for NFκB activation. To determine the role of NFκB in the increased cell death observed with *Lgr4*-deficiency in INS1 cells under basal and cytokine treatment, we used the NFκB inhibitor ammonium pyrrolidinedithiocarbamate (PDTC), which blocks NFκB activity (35–37). Indeed, the NFκB inhibitor significantly decreased β-cell death in *Lgr4*-deficient INS1 cells under basal conditions (Fig. 4G), validating the role of the RANK-TRAF6-NFκB pathway in the increased cell death observed with LGR4 deficiency. As expected, cytokines induced a ∼15-fold increase in cell death in sc-siRNA transfected cells, and the NFκB inhibitor protected against this increased cell death (Fig. 4H). Importantly, the further significant 2-fold increase in cell death observed with cytokines in *Lgr4*-deficient versus SC-siRNA treated INS1 cells was completely blocked by the NFκB inhibitor (Fig. 4H). This implies that NFκB mediates the increased basal and cytokine-induced β-cell death observed with LGR4 deficiency. Thus, mechanistically, LGR4 mediates its pro-survival effects on β-cells through suppressing stimulation of RANK-TRAF6 interaction and subsequent activation of the NFκB pathway.

### Young adult (10-13-week-old) *Lgr4*cko mice exhibit normal glucose metabolism and plasma insulin levels

Conditional β-cell-specific knockout (cko) of *Lgr4* using *Lgr4*-floxed (38, 39) and Ins1-*Cre*-knockin mice (40), resulted in a 50 to 75% reduction in *Lgr4* mRNA in islets of male (Fig. S1A) and female (Fig. S1B) *Lgr4*cko mice, when compared to wild-type (WT) controls, respectively. Body weight (Fig. S1, C, D) and random-fed blood glucose (Fig. S1, E, F) in male (4-10 weeks) and female (4-12 weeks) cko mice did not significantly differ from their WT littermates. Similarly, insulin tolerance test (ITT) (Fig. S1, G, H), intraperitoneal glucose tolerance test (IPGTT) (Fig. S1, I, J), or plasma insulin levels (Fig. S1, K, L) at 10-15 weeks of age showed no significant differences between *Lgr4*cko and WT control mice in males or females, suggesting that a chronic deletion of *Lgr4* in β-cells does not impair glucose metabolism in early adulthood.

### *Lgr4*cko mice exhibit impaired β-cell health when compared to WT controls under basal conditions as well as with increased metabolic stress

We next assessed the effects of *Lgr4*cko on β-cell health *in vivo*. As observed with acute knockdown of *Lgr4* in mouse islets *in vitro* (Fig. 2), β-cell proliferation assessed by pHH3-insulin co-staining (Fig. 5A) was significantly decreased under basal conditions in *Lgr4*cko female (Fig. 5B) but not male (Fig. 5C) mice, relative to WT control mice at 10-13 weeks of age. β-cell death measured by TUNEL-insulin co-staining (Fig. 5D), was significantly increased in both female (Fig. 5E) and male (Fig. 5F) *Lgr4*cko mice under basal conditions compared to controls, similar to our *in vitro* observations.

**Figure 5.**
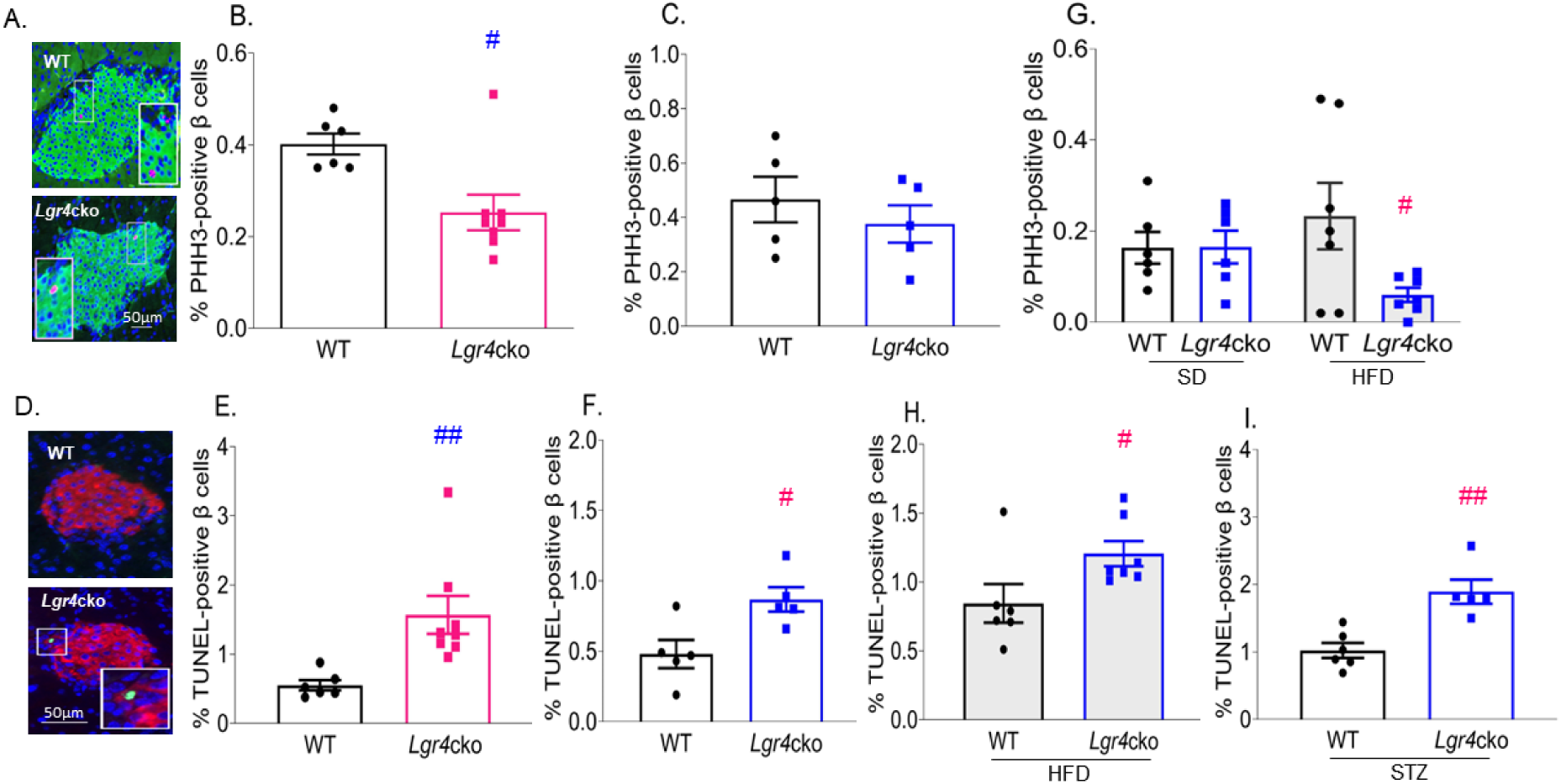
Young adult *Lgr4*cko mice exhibit impaired β-cell health compared to WT controls under basal and increased metabolic stress. (A) Representative images of immunofluorescent staining of mouse pancreatic sections from 13-week-old WT and *Lgr4*cko female mice for pHH3 (red), insulin (green), and DAPI (blue). Percent pHH3-positive β-cells in WT and *Lgr4*cko **(B)** 13-week-old female (n=6-8) and **(C)** 10-week-old male (n=5) mice. **(D)** Representative images of immunofluorescent staining of mouse pancreatic sections from 11-week-old WT and *Lgr4*cko male mice for TUNEL (green), insulin (red), and DAPI (blue). Percent TUNEL-positive β-cells in WT and *Lgr4*cko **(E)** 13-week-old female (n=6-8) and **(F)** 11-week-old male (n=5) mice. **(G)** Percent pHH3-positive β-cells and **(H)** percent TUNEL-positive β-cells in WT and *Lgr4*cko male mice fed SD or HFD (n=6-7) for 14 weeks. **(I)** Percent TUNEL-positive β-cells in WT and *Lgr4*cko male mice (n=5-6) at day 7 after 5 days of STZ treatment. ^#^p<0.05, ^##^p<0.01 vs the WT group with the same treatment. White bar indicates the scale for the immunofluorescent images (50µm). Individual symbols in the graphs represent individual mice. All data represent mean ± SEM. Statistical analysis was by t-test for comparison of two groups, and by ANOVA with Tukey’s post-hoc analysis for comparison of more than two groups.

Since *Lgr4*cko male mice, unlike female mice, did not exhibit reduced β-cell replication basally, we exposed them to increased metabolic demand through a high fat diet (HFD). *Lgr4*cko and WT male mice (10-weeks of age) were fed a HFD or standard diet (SD) for 14 weeks, with regular monitoring of body weight and blood glucose, along with an ITT and IPGTT at 23-24 weeks of age, followed by plasma insulin measurement and pancreas collection at the end of the study (Fig. S2A). Both WT and *Lgr4*cko mice on HFD showed a significant increase in body weight compared to WT and *Lgr4*cko mice on SD, respectively, starting at 12-weeks of age and continuing up to the end of the study (Fig. S2B). Similarly, the blood glucose in WT and *Lgr4*cko HFD mice started to significantly increase at 16-18 weeks of age compared to their counterparts on SD (Fig. S2C). However, there was no significant difference in body weight or blood glucose between WT/HFD and *Lgr4*cko/HFD mice (Fig. S2, B, C). Insulin resistance, as measured by ITT (Fig. S2D) and area under the curve (AUC) (Fig. S2E), was significantly increased in both WT and *Lgr4*cko mice on HFD compared to their respective controls on SD. Glucose tolerance as assessed by IPGTT (Fig. S2F) and AUC (Fig. S2G), showed impairment in the HFD versus SD groups, but was significant only in *Lgr4*cko/HFD versus *Lgr4*cko/SD mice, and not in WT/HFD versus WT/SD mice. Interestingly, β-cell homeostasis evaluated at 24-weeks of age showed a significant decrease in β-cell proliferation in *Lgr4*cko/HFD compared to WT/HFD mice (Fig, 5G). However, no significant change in proliferation was observed between *Lgr4*cko and WT mice under control SD even at 24 weeks of age (Fig, 5G), similar to what was observed basally in younger 10-week-old male *Lgr4*cko mice (Fig, 5C). β-cell death was also significantly increased in *Lgr4*cko/HFD compared to WT/HFD mice (Fig, 5H). We then examined the effect of multiple low dose streptozotocin (STZ) (MLDS), a model of β-cell destruction and inflammation, on β-cell health in male *Lgr4*cko mice. We used only male mice for these studies due to the inconsistent response of female mice to STZ (41, 42). 10-week-old male WT or *Lgr4*cko mice were injected with vehicle (Veh) or low-dose STZ for five consecutive days with body weight and blood glucose monitored daily and pancreas harvested at day 7 (Fig. S2H). There were no significant differences in body weight (Fig. S2I) or blood glucose (Fig. S2J) between the four groups of mice during the 7 days. As expected, β-cell death was significantly increased in the WT/STZ (Fig. 5I) compared to WT/Veh treated mice (Fig. 5F). Importantly, there was a further significant increase in β-cell death in *Lgr4*cko/STZ (1.89±0.18%) versus WT/STZ (1.02±0.11%) treated mice (Fig. 5I). These findings demonstrate a further impairment in β-cell health when stressors are combined with LGR4 deficiency.

### β-cell health deteriorates with aging in both males and females, but glucose homeostasis is impaired only in female Lgr4cko mice

Analysis of β-cell homeostasis in aged (23-month-old) *Lgr4*cko mice revealed a significant decrease in β-cell proliferation in female (0.08±0.02%) (Fig. 6A) as well as male (0.06±0.01%) (Fig. 6B) *Lgr4*cko mice compared to WT female (0.21±0.03%) and male (0.22±0.06%) mice. Unlike female *Lgr4*cko mice which show reduced β-cell replication through their adult life, male *Lgr4*cko mice display impaired proliferation only with aging or additional stressors. β-cell death was significantly increased in female (0.81±0.12%) (Fig. 6C) and male (0.27±0.05%) (Fig. 6D) *Lgr4*cko mice compared to their respective WT controls (0.24±0.08% and 0.11±0.05%, respectively). β-cell mass, quantified on insulin-stained pancreatic sections of female (Fig. 6E) and male (Fig. 6F) *Lgr4*cko and WT mice, was not significantly different in either female (Fig. 6G) or male (Fig. 6H) *Lgr4*cko versus WT control mice.

**Figure 6.**
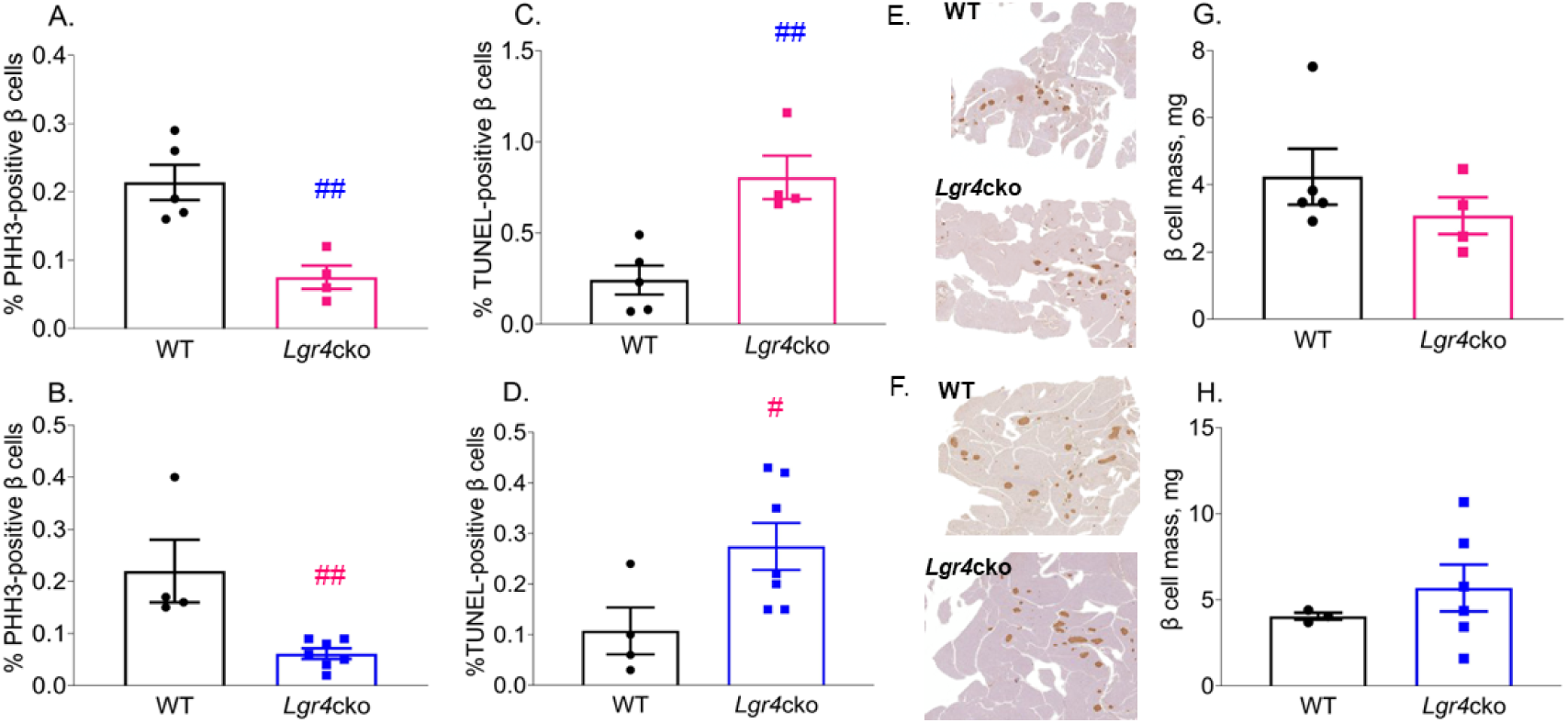
Aged *Lgr4*cko male and female mice exhibit reduced β-cell proliferation and increased β-cell death. β-cell histomorphometry in 23-month-old WT and *Lgr4*cko mice assessing for percent pHH3-positive β-cells in **(A)** female (n=4-5) and **(B)** male (n=4-7) mice; percent TUNEL-positive β-cells in **(C)** female (n=4-5) and **(D)** male (n=4-7) mice; immunohistochemical staining for insulin in **(E)** female and **(F)** male mice (representative images); β-cell mass in **(G)** female (n=4-5) and **(H)** male (n=3-6) mice. ^#^p<0.05, ^##^p<0.01 vs the WT group. Individual symbols in the graphs represent individual mice. All data represent mean ± SEM. Statistical analysis was done by t-test for comparison of two groups.

To assess the effect of aging on glucose homeostasis in *Lgr4*cko mice we measured body weight and blood glucose every month, starting at 8 months of age, and performed metabolic phenotyping at 11.5, 19.5, and 22.5 months of age in WT and *Lgr4*cko male and female mice. There was no significant difference in body weight between male WT and *Lgr4*cko mice (Fig. S3A). However, the female *Lgr4*cko mice had decreased body weight starting at 14-16 months of age, which remained significantly lower at the end of the study at 22-23 months of age relative to WT female mice (Fig. S3B). Blood glucose between WT and *Lgr4*cko mice was not significantly different in males (Fig. S3C) or females, except for a small but significant increase observed in female *Lgr4*cko mice at 12-14 months of age (Fig. S3D). ITT and its AUC assessed at 11-12-months (Fig. S3, E-H) or 19-20-months of age (Fig. S3, I-L) were not significantly different between WT and *Lgr4*cko mice in either males or females. The IPGTT response and AUC were similar in control WT and *Lgr4*cko male mice at 11.5 (Fig. 7, A, B) and 19.5 (Fig. 7, C, D) months of age. However, female *Lgr4*cko mice exhibited significantly impaired glucose tolerance and AUC compared to WT littermates at 11.5 (Fig. 7, E, F) and 19.5 (Fig. 7, G, H) months of age. Plasma insulin measured in response to the glucose challenge, although significantly increased at 15-min compared to the fasting (0-min) time-point in both WT and *Lgr4*cko male mice (Fig. 7I), was similar between *Lgr4*cko and WT male mice at both time-points (Fig. 7I). However, female *Lgr4*cko mice had significantly reduced plasma insulin levels during fasting and 15-min post-glucose stimulation, compared to WT controls (Fig. 7J). Unlike the WT female mice, which had a significant increase in plasma insulin levels 15 minutes after a glucose challenge, *Lgr4*cko female mice did not (Fig. 7J). Likewise, endpoint plasma insulin levels at 23 months of age were similar in male *Lgr4*cko and WT mice (Fig. 7K), but female *Lgr4*cko mice had significantly 3-fold lower plasma insulin levels (1.06±0.34µg/l) relative to their WT littermates (3.04±0.40µg/l) (Fig. 7L).

**Figure 7.**
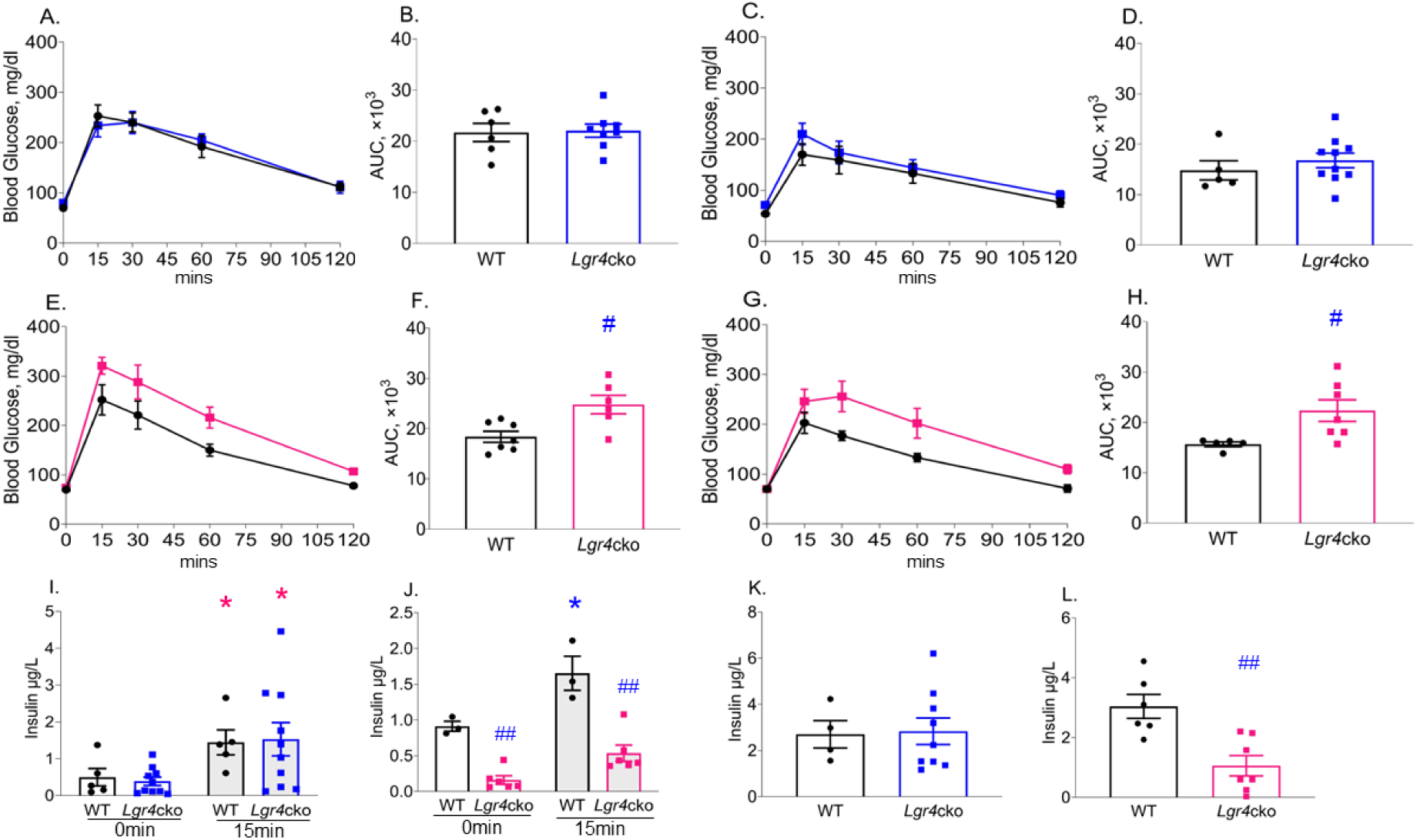
Aging impairs glucose homeostasis in female but not male Lgr4cko mice. Glucose clearance (A, C, E, G) and area under the curve (B, D, F, H) during IPGTT in aged WT (black symbols) and **(A-D)** *Lgr4*cko male (blue symbols) mice at (A, B) 11-12 months (n=6-8) and (C, D) 19-20 months (n=5-10) of age, and **(E-H)** Lgr4cko female (pink symbols) mice at (E, F) 11-12 months (n=6-7) and (G, H) 19-20 months (n=5-7) of age. **(I, J)** Plasma insulin in WT and *Lgr4*cko mice at 0- and 15-min time points of the IPGTT performed at 19-20 months of age in (I) male (n=5-10) and (J) female (n=3-6) mice. **(K, L)** Plasma insulin in WT and *Lgr4*cko mice at 23 months of age in (K) male (n=4-9) and (L) female (n=6-7) mice. ^#^p<0.05, ^##^p<0.01 vs the WT group at the same time-point. *p<0.05 vs 0min in the same group. Individual symbols in the bar graphs represent individual mice. All data represent mean ± SEM. Statistical analysis was by t-test for comparison of two groups, and by ANOVA with Tukey’s post-hoc analysis for comparison of more than two groups.

In summary, *Lgr4* deficiency in β-cells impairs β-cell health, increasing β-cell death in young adult and aged male and female mice, and reducing β-cell proliferation, but with a more severe impact in female mice where replication is reduced in young and aged adults, versus male mice, where impairment in proliferation occurs only with aging and stress. Glucose homeostasis is normal in young adult *Lgr4*cko male and female mice. However, with aging impaired glucose tolerance and reduced plasma insulin are observed in female but not male *Lgr4*cko mice.

## Discussion

LGR4 is expressed in rodent and human β-cells, and despite being the fourth most abundant GPCR in human islets (9), its role in this endocrine cell population remains unknown. In this study we demonstrate that LGR4 is vital for maintaining β-cell homeostasis through regulation of β-cell proliferation and survival, in basal and stress conditions, *in vitro* and *in vivo* and is negatively regulated by multiple stressors related to T1D and T2D in rodent and human islets. We have leveraged *Lgr4* loss and gain of function studies to demonstrate that LGR4 not only protects β-cells against multiple stressors but is essential for basal β-cell survival. Using multiple approaches, we demonstrate that LGR4/RANK stoichiometry is critical for β-cell survival, and that LGR4 protects β-cells through the suppression of RANK/TRAF6 interaction and subsequent activation of the NFκB pathway.

Our data show that *Lgr4* mRNA levels are reduced in islets in response to acute (treatment with proinflammatory cytokines), or chronic (increased metabolic demand in the HFD fed mice, db/db mice, or with aging) stress in both mice and humans. Our analysis of the published bulk RNAseq data sets indicated a decrease in *LGR4* mRNA in human islets treated with IL1β and IFNγ (43), or with Brefeldin A, a stress inducer of the Golgi Apparatus (44). Reduction of *LGR4* has been observed in multiple tissues under varied stressors. A proinflammatory milieu associated with venous ulcer formation negatively affected *LGR4* mRNA levels (24). Women with gestational diabetes have reduced *LGR4* in the placenta compared to healthy subjects (45), and plasma LGR4 levels decline with progression of hypertension in T2D (46).

Using two independent methods to acutely knockdown *Lgr4* in rodent β-cells *in vitro* (*Lgr4*-siRNA and *Lgr4*-floxed mouse islets transduced with Ad-*Cre*), we show that LGR4 plays a crucial role in pancreatic β-cell survival in male and female mouse islets. Our data also establish a protective effect of LGR4 overexpression on primary rodent and human β-cell survival under cytokine-induced stress. *Lgr4* deficiency causes inflammation and cell death in many other tissues. In particular, *LGR4* knockdown increases TNFα-mediated primary hepatocyte cell death in culture and in the setting of stress conditions-hepatic ischemia/reperfusion-induced injury (47) and lipopolysaccharide/D-Galactosidase-induced liver injury (48). *Lgr4* knockout also leads to a more pronounced inflammatory bowel disease under dextran sodium sulfate administration (21); results in increased cell death in the renal peripheral mesenchyme, accompanied by decreased levels of anti-apoptotic protein PAX2 (22); and manifests with impaired skin regeneration leading to venous ulcer formation in mice and humans (24). On the other hand, overexpression of LGR4 in osteoarthritic synoviocytes results in decreased pro-inflammatory cytokine production and inhibition of NF-κB activity, thus ameliorating osteoarthritic inflammation (23). Taken together, these studies along with our data indicate the importance of LGR4 as an anti-inflammatory pathway, with the potential to leverage it for diabetes-related β-cell stress prevention.

Our mechanistic studies identified an intriguing link between the LGR4 and RANK receptors in β-cells. A connection between these two pathways was first recognized in osteoclasts, through identification of a common ligand, RANKL, that can activate both receptors, but produce opposite effects. LGR4 overexpression in osteoclast precursor cells (RAW264.7) decreased RANK-TRAF6 interaction, leading to reduced NFκB pathway activation (15). Prior work from our group has shown that in rodent and human β-cells the RANKL/RANK pathway is essential for mediating cytokine-induced cell death through its interaction with the TRAF6 adaptor protein leading to activation of the NFκB pathway (20). We now show that the pro-survival effect of LGR4 on β-cells is mediated through the suppression of RANK/TRAF6 interaction and inhibition of NFκB activation. The increase in basal β-cell death observed with LGR4 deficiency was reversed upon simultaneous deletion of the RANK receptor, or by specifically inhibiting RANK/TRAF6 interaction using a peptide inhibitor. Furthermore, we found that NFκB was activated in *Lgr4*ko β-cells and inhibiting this pathway could protect *Lgr4*ko cells from basal and cytokine-induced cell death. Similar to our results, *Lgr4*ko in keratinocytes leads to decreased phosphorylation of p65 at Ser468, leading to NFκB-mediated inflammation and impaired skin wound healing (24). Thus, LGR4 protects β-cells by inhibiting RANK/TRAF6 interaction and suppressing subsequent NFκB activation.

In the context of LGR4 ligands, previous studies (49–51) have shown contradictory effects of RSPO1, a classical LGR4 ligand, on β-cell health. Wong et al. (49) demonstrated that RSPO1 acted as a mitogen for pancreatic β-cells (MIN6 cell line and primary mouse islets), protected them against proinflammatory cytokine-induced cell death, and stimulated insulin secretion *in vitro*. Surprisingly, a subsequent study by the same group (50) performed on whole body *Rspo1* knockout mice showed improved β-cell mass and better glycemic control compared to control mice. STZ injection in the *Rspo1* knockout mice resulted in a lower number of apoptotic β-cells, increased β-cell neogenesis and maturation compared to the WT controls (51). Based on these contradictory *in vitro* and *in vivo* findings, the role of the LGR4 ligand, RSPO1, on β-cells remains unclear. Our data showing that *Rank*ko could rescue the increased death observed in *Lgr4*ko INS1 cells implies that RSPO1/LGR4 interaction is unlikely to mediate the pro-survival effects of LGR4 on β-cells, but rather that RANK/LGR4 stoichiometry determines the effect on β-cell survival.

Our study demonstrates that LGR4 also regulates β-cell proliferation. However, unlike β-cell survival, which is modulated in both male and female mouse islets, we observed sex-specific effects of LGR4 on β-cell replication, where β-cells from female, but not male, mice had significantly reduced proliferation upon *Lgr4* knockdown under basal conditions *in vitro*. Our *in vivo* studies in young adult β-cell specific *Lgr4*cko mice showed a similar phenotype of reduced β-cell proliferation only in female mice under basal conditions, as observed *in vitro*. However, when male Lgr4cko mice were subjected to 14-weeks of HFD, a stressor related to T2D, they showed a significant reduction in β-cell replication compared to control WT mice. The sex-specific difference observed in β-cell proliferation in young adult *Lgr4*cko mice could be due to impairment in estrogen signaling under *Lgr4*cko. RSPO1/LGR4 upregulates estrogen receptor ɑ (ERɑ) expression in mammary luminal cells independent of the Wnt signaling pathway (52). ERɑ drives β-cell replication during development in the mouse pancreas (53), as well as *in vitro* in rodent and human β-cells (54). Future studies will determine whether LGR4 deficiency induces sex-specific differences in β-cell proliferation through the regulation of ERɑ.

As seen *in vitro*, *Lgr4* deficiency *in vivo* increased β-cell death basally in both male and female mice, and further exacerbated β-cell death under stressors such as HFD, as well as with MLDS-treatment, a stressor related to T1D. Interestingly, the changes in β-cell proliferation and/or survival under basal conditions did not result in aberrant glucose homeostasis in young adult *Lgr4*cko mice. Upon aging, both male and female *Lgr4*cko mice showed a significant reduction in β-cell proliferation and survival, without any significant effect on β-cell mass, relative to WT control mice. However, only female aged *Lgr4*cko mice displayed glucose intolerance and lower plasma insulin levels during the IPGTT and at the end of the study. This is likely due to a combination of the persistent life-long reduction in β-cell proliferation and survival seen in these mice; and likely from reduced β-cell function as evidenced by reduced plasma insulin levels despite normal β-cell mass.

Our studies identify LGR4, a GPCR, as a crucial regulator of β-cell proliferation and survival *in vivo* under basal and stress-induced conditions. GPCRs are key mediators of essential cellular processes and therapeutic targets for numerous diseases, including the GLP1R for diabetes (7). Conditional β-cell specific knockout of the *Glp1r* eliminates insulin responses to intravenous and intraperitoneal GLP1 administration (55). Barrella et al. demonstrated that conditional β-cell specific knockout of another member of the GPCRs, *β-arrestin-1* (*barr1*), leads to impaired β-cell replication and function in insulin-resistant mice maintained on an obesogenic diet (56). However, thus far, we do not know of any GPCRs that regulate β-cell expansion and survival, basally *in vivo*, further underscoring the significance of our findings demonstrating the importance of LGR4 in preserving normal β-cell health and homeostasis. Our studies highlight the importance of maintaining a proper stoichiometry between the two receptors, LGR4, a positive, and RANK, a negative (20), modulator for β-cell survival. The pancreatic β-cell, a central player in metabolic regulation, can be added to the repertoire of cell types regulated by LGR4. Future studies will evaluate the therapeutic implications of this receptor and its potential ligand(s) in diabetes.

## Materials and Methods

### Animal studies

For *in vitro* studies, islets were isolated from both male and female mice from the following strains and age: for effects of *Lgr4* knockout on β-cell proliferation and death assays islets from 10-12 (β-cell proliferation) or 13-24 (β-cell death) week-old *Lgr4*-floxed mixed background mice (kindly shared by Dr. Katsuhiko Nishimori, Tohoku University, Japan) were transduced with Adv-*LacZ* or Adv-*Cre*. To assess the effects of *Lgr4* overexpression on β-cell death, 16-24-week-old C57BL/6 male and female mice (Jackson Laboratories, Bar Harbor, ME) were used. For *in vivo* studies, female and male WT (*Lgr4*fl/fl-Ins1*Cre*-negative) and *Lgr4*cko (*Lgr4*fl/fl-Ins1*Cre*-positive) mice were used. For basal phenotyping, body weight and blood glucose were measured once a week in young 4-10-week-old males and 4-13-week-old females, or once a month in aged 8-22-month-old males and females. Blood and pancreata samples were harvested at 12 weeks of age in young females, and 22-23 months of age in aged males and females. For the HFD studies, body weight and blood glucose were measured weekly on 10-week-old male WT and *Lgr4*cko mice fed regular chow (SD) or 60% kcal HFD (Research Diets, NY) for 14 consecutive weeks. Blood and pancreata samples were harvested at 24 weeks of age. For MLDS studies, 10-week-old male WT and *Lgr4*cko mice were injected with vehicle (sodium citrate, pH 4-4.5) or low-dose (40mg/kg body weight) (19, 57, 58) STZ (Sigma Aldrich, St Louis, MO) for five consecutive days. Body weight and blood glucose were measured daily for seven days and pancreata and serum were harvested at 11-weeks of age. Animal studies were performed with the approval of, and in accordance with, guidelines established at the Beckman Research Institute, City of Hope, Duarte, CA, and principles of laboratory animal care were followed.

### Mouse and human islet isolation and cell culture: Adv-transduction

Mouse islets (from *Lgr4*-floxed or C57Bl6/J mice) were isolated by collagenase digestion and histopaque gradient separation and cultured in complete medium [RPMI supplemented with 5.5mM glucose, 10% FCS, and 1% penicillin-streptomycin (pen/strep)], as described previously (20, 59). Human islets (Supplemental Table 1) were obtained from the Integrated Islet Distribution Program (IIDP), Prodo Labs, and Southern California Islet Resource Center. Rodent and human islets were cultured in complete media for at least 24h before they were hand-picked under the microscope for treatment and analysis. Islet cell cultures were prepared by 10-min trypsinization with intermittent pipetting. Cells from 50 islet equivalents (IEQs; 1 IEQ=125 μm diameter) were plated on coverslips in a 24-well plate initially in a small volume (50 μl) of media for 2h to allow attachment. To generate *Lgr4*ko *in vitro* acutely in islet cells, trypsinized *Lgr4*-floxed islet cell cultures were transduced with recombinant Adv-*LacZ* (Ctrl) or Adv-*Cre* recombinase (Gene Transfer Vector Core, Iowa City, IA). For overexpression studies, C57Bl6/J mouse or human islet cell cultures were transduced with recombinant Adv-*Cre* (as control) or Adv-*Lgr4*. In all experiments, recombinant adenoviruses were transduced at a multiplicity of infection (moi) of 100 in a volume of 50 μl of complete media for 2h, after which 1 ml of complete islet media was added. After 48h or 72h cells were fixed in 2% paraformaldehyde for 30 min, to stain for Cre recombinase (1:400, Cell Signaling Technology, Denvers, MA) or TUNEL (Promega, Indianapolis, IN), respectively; or harvested for RNA analysis after 72h (20).

### Assessment of β-cell proliferation and death *in vitro* in dispersed mouse or human islet cells

For basal and cytokine-treated cell death experiments, islet cells were cultured in serum-containing medium and a mix of species-compatible pro-inflammatory cytokines, IL1β 50U (mouse, 0.091 ng/ml; human, 0.714 ng/ml), IFNγ 1000U (mouse, 118.62 ng/ml; human, 50 ng/ml), and TNFα 1000U (mouse, 3.7 ng/ml; human, 13.15 ng/ml) (R&D Systems, Minneapolis, MN) or vehicle (0.1% PBSA) was added for 16-24h. β-cell death was assessed by staining for insulin (1:800, Abcam, Cambridge, MA), TUNEL (Promega, Indianapolis, IN) and 4’,6-diamidino-2-phenylindole (DAPI) (Life Technologies), using Alexa Fluor 594 anti-insulin (Life Technologies, Carlsbad, CA) as a secondary antibody, and quantified as percentage of TUNEL-insulin to total insulin-positive cells (20, 59). β-cell proliferation was measured by insulin, pHH3 (1:1000, Sigma Millipore, Chicago, IL) and DAPI staining, and quantified as percentage of pHH3-insulin to total insulin-positive cells (19, 60). Reagents and antibodies used for the *in vitro* studies are listed in Supplemental Table 2. An average of at least 1776±311 and 1307±107 β-cells/condition for each sample were counted for the *in vitro* mouse and human islet studies, respectively.

### INS1 cell assays

Rat insulinoma cell line, INS1 and INS1 832/13, kindly shared by Drs. Patrick Fueger and Hirotake Komatsu (COH) was cultured in RPMI media supplemented with 11mM glucose, 10% FCS, 1% pen/strep, as described previously (20, 59, 61). Cells were regularly screened for the presence of mycoplasma. To knockdown *Lgr4*, INS1 cells seeded at 400×10^3^ cells/well in a 6-well plate for 24h, were cultured in pen/strep-free media overnight and transfected with 25 nmol/L of either *Lgr4* (Thermo Fisher Scientific) or scrambled (SC) Control No. 1 siRNA, (Thermo Fisher Scientific) siRNA using Lipofectamine RNAiMAX (Thermo Fisher Scientific) (61, 62). For *Lgr4* overexpression, INS1 cells seeded at 250×10^3^ cells/well in a 6-well plate were transduced with 100 moi of Ad-*Cre* (control) or Ad-*Lgr4* for 48h (60, 61). Un-transfected or un-transduced, *Lgr4* and SC siRNA-transfected, or Ad-*Cre* and Ad-*Lgr4*-transduced INS1cells were collected for RNA 48h post-transduction or plated at a density of 50×10^3^ cells/well on coverslips in a 24-well plate to assess cell proliferation or death. For proliferation assay, un-transfected, SC or *Lgr4* siRNA-transfected INS1 cells treated with 10µM/ml Bromodeoxyuridine (BrdU) (Thermo Fisher Scientific, Milwaukee, WI) for 2h were stained for BrdU with a primary anti BrdU antibody (1:500, Abcam, Cambridge, MA). BrdU-positive cells were visualized using Alexa Fluor 594 anti-rat secondary antibody (Life Technologies) and DAPI (Life Technologies) as described earlier (63). β-cell proliferation was quantified in ImageJ program (National Institute of Health) (64), as percentage of BrdU-positive cells over total DAPI-positive cells (65), with an average of 7339±266 INS1 cells counted/well.

For cell death assay, un-transfected or un-transduced, *Lgr4* and SC siRNA-transfected, or Ad-*Cre* and Ad-*Lgr4*-transduced INS1cells were treated with vehicle (0.1% PBSA) or proinflammatory cytokines (IL1β, TNFα, and IFNγ) at concentrations described in Supplementary Table 2 for 16 or 24h and fixed in 2% PFA for 30 mins (20). For NFκB inhibition assay, *Lgr4* and SC siRNA transfected INS1 cells were seeded at a density of 50×10^3^ cells/well for 48h, and preincubated with PDTC or vehicle (DMSO, Fisher Scientific, Pittsburgh, PA) for 30 mins before addition of cytokines. To assess the role of the RANK pathway in *Lgr4*-deficient INS1 cell death, INS1 cells were transfected with SC, *Lgr4*, *Rank* (125, 25 nmol and 100nmol/L, respectively), or *Lgr4*+*Rank* siRNAs and RNA expression as well as basal and cytokine-induced cell death was assessed as described above. To test the role of RANK-TRAF6 interaction we pre-treated INS1 cells seeded for 48h at a density of 50×10^3^ cells/well with 30 µM of the TRAF6 inhibitor peptide (Novus Biologicals, CO) for 30 min before treating with the cytokine mix. Cleaved-Casapase-3 staining was performed using anti-Cleaved-Casapase-3 antibody (1:200, Cell Signaling Technologies, MA) and secondary Alexa Fluor 488 anti-rabbit antibody (Life Technologies) and DAPI. Cell death was quantified as percentage of Cleaved-Casapase-3-positive cells over total DAPI-positive cells (59, 62), with an average of 7096±332 INS-1 cells counted/well. Phospho p65 Ser468 levels were assessed by staining for phospho p65 Ser468 (1:200; Thermo Fischer Scientific, Waltham, MA) using secondary Alexa Fluor 488 anti-rabbit antibody and DAPI. phospho p65 Ser468 levels were quantified as percentage of phospho p65 Ser468-positive nuclei over total DAPI-positive cells, using Image J, with an average of 5595±589 INS-1 cells counted/well. For all *in vitro* studies, samples were analyzed in duplicate in a blinded fashion.

### RNA Analysis

RNA was isolated using the RNeasy Kit (#74004; Qiagen, Hilden, Germany) from INS1 cells (400,000 cells/sample); from whole mouse islets (100–300 IEQ/animal); and from mouse and human islet cell cultures (100-150×10^3^ cells/well). cDNA was synthesized using either the qScript SuperMix (Quanta Bio, Beverly, MA) or SuperScript III reverse transcription kit (Life Technologies) (19, 20). Gene expression was analyzed by quantitative real-time (qRT) PCR using specific primers and Taqman probes (Supplementary Table 2).

### Glucose homeostasis

Blood glucose was measured on tail snips using a portable glucometer (Freestyle Lite, Amazon). ITT and IPGTT were performed at 11-12 weeks of age in young mice under basal conditions, at 23-24 weeks of age in the HFD study, at 11-12, 19-20, and 22-23 months of age in the aging study. Before performing IPGTT, mice were fasted for 16-18h and injected with 2g glucose/kg body weight. Blood glucose was measured pre- and at regular intervals starting at 15 to 120 mins post-glucose treatment. Plasma insulin was measured on blood drawn at fasting, after 15 min of glucose challenge during IPGTT, and before termination of the study, using an insulin ELISA kit (Mercodia, Inc, Winston Salem, NC) (20, 59). For ITT, the mice were injected in the afternoon with human insulin (Humulin; Eli Lilly, Indianapolis, IN) 1.5U/gm body weight, and blood glucose was measured pre- and at regular intervals starting at 15 to 120 mins post-insulin treatment (59).

### Assessment of β-cell proliferation, death, and mass in pancreatic sections *in vivo*

Pancreata were weighed, fixed in 10% neutral buffered formalin (Sigma, St Louis, MO), and paraffin embedded. Pancreatic sections were stained with antibodies against insulin (1:800) and pHH3 (1:500), after antigen retrieval with high pressure cooker steam in citrate buffer for 20 min, using an immunofluorescence secondary antibody, as indicated in Supplemental Table 2. β-cell proliferation was quantified as percentage of pHH3-insulin to total insulin-positive cells. β-cell death was assessed on pancreatic sections stained with antibodies against insulin and TUNEL, visualized under a fluorescent microscope (Zeiss Observer, San Diego, CA) and quantified as percentage of TUNEL-insulin to total insulin-positive cells. For β-cell mass assessment, mouse pancreatic sections were incubated overnight at 4°C in guinea pig anti-insulin antibody (1:800) and visualized using the diaminobenzidine peroxidase substrate (Vector laboratories, Burlingame, CA) and hematoxylin staining. Histomorphometry for β-cell mass was performed in a blinded way on 4-6 insulin-stained pancreatic sections per animal separated by 50μm each, using the Image J program (66). β-cell mass was quantified per animal as the ratio of the insulin-positive to total pancreatic area, multiplied by the pancreas weight and averaged for all the sections/mouse (59).

### Omic data analysis for *Lgr4* expression

*Lgr4* mRNA levels in mouse islets were retrieved from the single cell RNA-analysis based published data set (26), available at: https://tabula-muris.ds.czbiohub.org/. The differential gene-expression data for the db/db and HFD mouse models is a meta-analysis of publicly available datasets. The analysis using the db/db model is based on the RNAseq data (GSE132261) on islets isolated from 4-week-old db/db mice and control BKS littermates that were reported by Jaafar et al. (28). The differential gene expressions (DEG) list was generated as per methods reported in the paper and was made accessible to us by the corresponding author Dr. A. Bhushan. The analysis of the islets in the context of a HFD is based on the RNA-seq data by Wang et al. (27) that compared islets from mice fed with a chow diet or HFD for a period of two months (HFD_Time_Scale. csv, hosted at https://data.mendeley.com/datasets/mwgxv7m927/2). Heatmaps were generated using the Morpheus utility (https://software.broadinstitute.org/morpheus/; RRID:SCR_017386). Each heatmap row in the db/db dataset represents log transformed normalized read counts. Color scale is relative to the range of values in each row. The rows in the HFD heatmaps represent normalized read counts. Color scale is relative to the range of values in each row.

We measured the expression of *LGR4* in single-cell (sc) and single-nucleus (sn) RNA sequencing data on human islets obtained from a previous publication (29). Briefly, we performed donor-matched scRNA and snRNA sequencing preparation. We aligned reads using the CellRanger pipeline (V6.1.1), created count data with the Seurat package (V4.1.1) after quality control, and integrated each sample with the SCT integration algorithm. Expression data were obtained from the RNA assay and visualized. Bulk RNAseq analysis was performed on control non-diabetic human islets (150-200IEQ) (Supplemental Table 1) treated for 24h with vehicle (0.1% PBSA) or a mix of human cytokines, as described above. The RNA-seq fastq reads were processed using Partek Flow Genomics pipeline version 10.0 (67). Raw fastq reads were preprocessed to determine low quality reads and contaminants using the Partek QA/QC module, followed by trimming bases at both ends based on a minimum Phred score of 30 using the trim bases module. The cleaned reads were aligned to the reference genome (hg38) using STAR version 2.7.8a (68). Gene counts were quantified using a modified form of the Expectation/Maximization algorithm (Partek E/M) which is part of Partek Genomics pipeline) (69). The raw gene counts were normalized in Partek based on sample library sizes and converted to counts per million (CPM) by adding a pseudocount of 10^-4^. The resulting normalized counts were batch corrected by fitting to a generalized linear model using the Partek Batch correction module, before calculating DGEs. P values of DGEs were estimated using the ANOVA module in Partek, followed by conversion to false discovery rates (FDR) according to the Benjamini-Hotchberg method (70). Heatmap was constructed using the ComplexHeatmap package (71) as part of the R software version 4.2 (72). An independent set of human islet preps (Supplemental Table 1) were used to confirm decrease in LGR4 mRNA under cytokine treatment vs. control by qPCR. Data for *LGR4* mRNA levels in EndoC-βH5 human β cell line was retrieved from bulk RNA analysis published by Frørup et al. (30), available at: https://www.ncbi.nlm.nih.gov/geo/geo2r/?acc=GSE218735. LGR4 expression values obtained under cytokine treatment for each experiment were normalized to vehicle control and expressed as average fold change.

## Statistical Analysis

Data are expressed as means ± SEM. Statistical significance was considered at p<0.05, p<0.01 p<0.001, and p<0.0001, for single, double, triple, and quadruple symbols, respectively, determined by unpaired two-tailed Student’s t-test (GraphPad Prism) for comparison between two groups; by One-way Analysis of Variance (ANOVA) with Tukey’s post-hoc HSD (http://astatsa.com/) for comparison between more than two groups.

## Abbreviation list

Ad: adenovirus
ANOVA: Analysis of Variance
AUC: Area Under the Curve
BrdU: Bromodeoxyuridine
CC3: Cleaved-Caspase-3
Cko: Conditional knockout
DAPI: 4′,6-diamidino-2-phenylindole
Fl: Floxed
GLP1R: Glucagon Like Peptide 1 Receptor
GPCR: G-protein coupled receptor
HFD: high fat diet
IPGTT: intraperitoneal glucose tolerance test
ITT: insulin tolerance test
ko: knockout
LGR4: Leucine Rich Repeat-containing G protein-coupled Receptor 4
MLDS: multiple low dose streptozotocin
moi: multiplicity of infection
NFKB: Nuclear factor kappa-light-chain-enhancer of activated B cells
pHH3: phospho-Histone H3
PTDC: Pyrrolidine dithiocarbamate
RANK: Receptor activator of NF-κB
RANKL: Receptor activator of NF-κB ligand
RSPO1: Rspondin 1
SC: Scramble silencing RNA
SD: Standard Diet
STZ: Streptozotocin
TNF: Tumor Necrosis Fact
TRAF6: Tumor necrosis factor receptor associated factor
T1D: Type 1 diabetes
T2D: Type 2 diabetes
Veh: vehicle
WT: wild-type

## Acknowledgments

We are grateful to the NIDDK-supported Integrated Islet Distribution Program, Southern California Islet Resource Center and Prodo Laboratories for providing human islets; to Rosemary Li, for performing qPCR analysis on the mouse islets treated with proinflammatory cytokines, to Dr. Navendu Vasavada for expertise on statistical analysis; to the microscopy cores at City of Hope (Dr. Brian Armstrong), and the pathology core at City of Hope supported by the National Cancer Institute of the National Institutes of Health under grant number P30CA033572, for assistance and use of their facilities; and to the donors of human islets.

## Author contributions

*Conceptualization:* J.F. and R.C.V. *Methodology:* J.F., P.W., R.K., G.L., Y.C.Y, S.B., S.D., A.G.O., N.G.K. and R.C.V. *Investigation:* J.F., Z.C., N.L.R., R.K., Y.C.Y, S.B., S.D., N.G.K. *Funding acquisition:* J.F., SD, N.G.K. and R.C.V. *Supervision:* R.C.V. *Writing-original draft: J*.F. and R.C.V. *Writing-review and editing:* J.F., Z.C., N.L.R., P.W., R.K., G.L., S.B., S.D., A.G.O., N.G.K., and R.C.V.

## Data and materials availability

All data needed to evaluate the conclusions in the paper are present in the paper and/or the Supplementary Materials.

## Supplementary Materials

**Figure S1.**
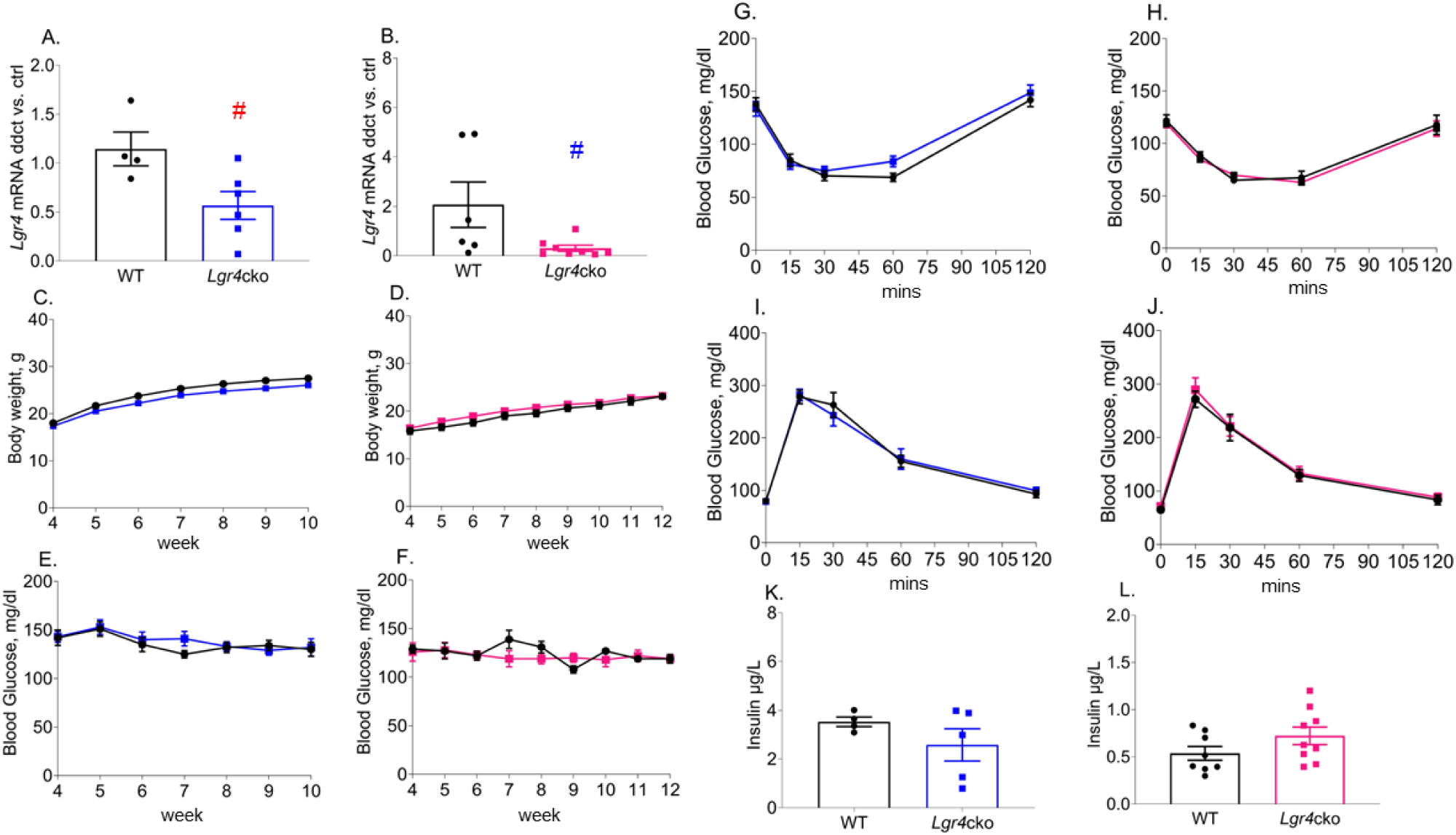
Young adult (10-13-week-old) *Lgr4*cko mice exhibit normal glucose metabolism and plasma insulin levels. qPCR analysis of *Lgr4* relative to *Cyclophilin A* or *Beta Actin* mRNA as housekeeping genes in islets from wildtype (WT) and *Lgr4*cko **(A)** 18-25-weeks old male (n=4-6) and **(B)** 13-weeks old female (n=6-7) mice. Phenotyping WT (black symbols) and *Lgr4*cko (male blue and female pink symbols, respectively) mice to assess **(C, D)** weekly body weight starting at 4 weeks of age in (C) male (n=11) and (D) female (n=6-8) mice; **(E, F)** weekly blood glucose starting at 4 weeks of age in (E) male (n=11) and (F) female (n=6-8) mice; **(G, H)** insulin tolerance in 8-9-week old (G) male (n=10) and (H) female (n=9-10) mice; **(I, J)** intraperitoneal glucose tolerance in 9-10-week old (I) male (n=10) and (J) female (n=9-10) mice; and **(K, L)** plasma insulin at 11-15 weeks of age in (G) male (n=4-5) and (H) female (n=8-9) mice. ^#^p<0.05 vs WT mice. Individual symbols in the bar graphs (A, B, K, L) represent individual mice. All data represent mean ± SEM. Statistical analysis was done by t-test.

**Figure S2.**
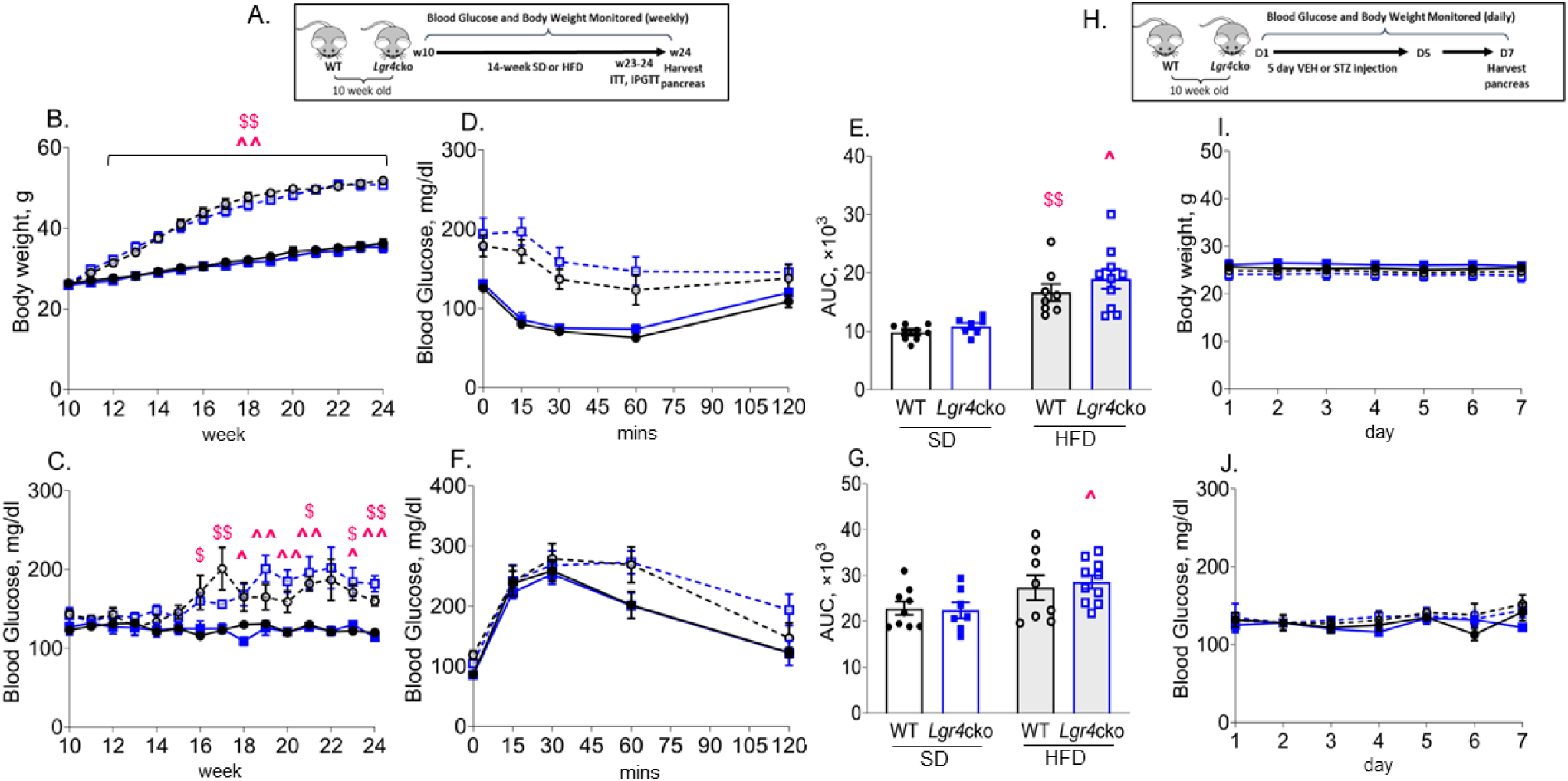
Glucose homeostasis in WT and *Lgr4*cko mice under HFD and MLDS. (A) Experimental design showing 10-week-old male wildtype (WT) and *Lgr4*cko mice fed a standard chow diet (SD) or a high fat diet (HFD) for 14 weeks with body weight and blood glucose measured weekly up to 24 weeks of age, ITT and IPGTT performed at 23-24 weeks of age, and pancreata harvested at 24 weeks at the end of the study. Phenotyping of these four groups of mice, WT/SD (filled black circles and line) (n=9), *Lgr4*cko/SD (filled blue squares and line) (n=7), WT/HFD (empty black circles, dashed black line) (n=8), and *Lgr4*cko/HFD (empty blue squares, dashed blue line) (n=10), to assess **(B)** weekly body weight from 10-24 weeks of age; **(C)** weekly blood glucose from 10-24 weeks of age; **(D)** ITT; **(E)** area under the curve for ITT; **(F)** IPGTT; and **(G)** AUC for IPGTT. **(H)** Experimental design showing 10-week-old male wildtype (WT) and *Lgr4*cko mice treated with vehicle (Veh) or low dose streptozotocin (STZ) daily for 5 days, with body weight and blood glucose measured daily up to day 7, and pancreata harvested at the end of the study at day 7. Phenotyping of these four groups of mice, WT/Veh (filled black circles and line) (n=5), *Lgr4*cko/Veh (filled blue squares and line) (n=5), WT/STZ (empty black circles, dashed black line) (n=6), and *Lgr4*cko/STZ (empty blue squares, dashed blue line) (n=5) mice to assess daily **(I)** body weight; and **(J)** blood glucose. ^$^p<0.05, ^$$^p<0.01 WT/HFD vs WT/SD mice; ^^^p<0.05, ^^^^p<0.01 *Lgr4*cko/HFD vs *Lgr4*cko/SD mice. Individual symbols in the bar graphs (E, G) represent individual mice. All data represent mean ± SEM. Statistical analysis was done by t-test.

**Figure S3.**
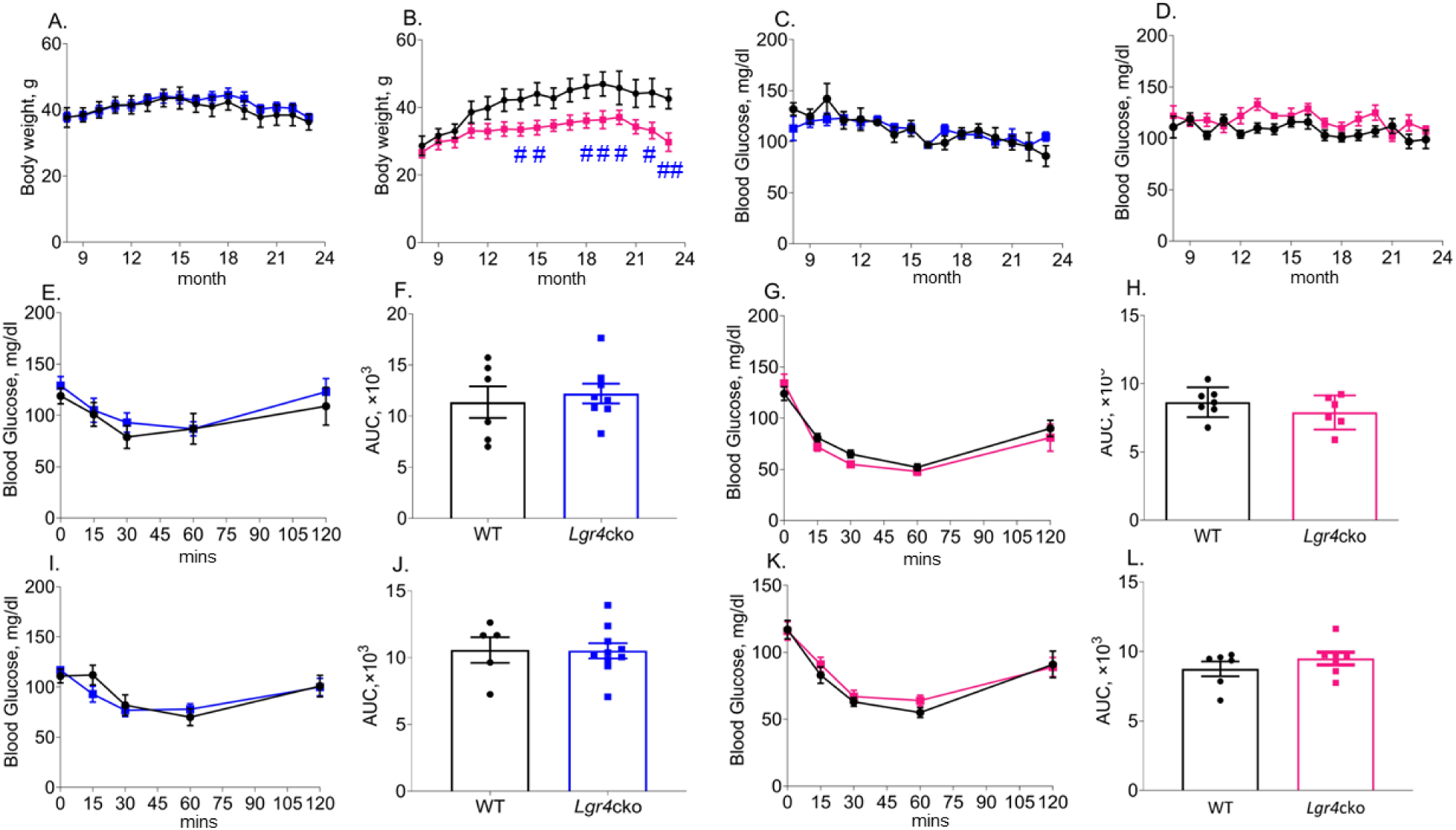
Glucose homeostasis in aged WT and *Lgr4*cko mice. Monthly assessment starting at 9 and up to 23 months of age in WT male and female (filled black circles and line), *Lgr4*cko male (filled blue squares and line), and *Lgr4*cko female (filled pink squares and line) mice, of body weight in **(A)** male (n=4-14) and **(B)** female (n=5-11) mice; blood glucose in **(C)** male (n=4-14) and **(D)** female (n=5-11) mice; ITT and AUC for ITT at 11-12 months of age in **(E, F)** male (n=4-14) and **(G, H)** female (n=5-11) mice; ITT and AUC for ITT at 19-20 months of age in **(I, J)** male (n=5-10) and **(K, L)** female (n=5-7) mice. ^#^p<0.05, ^##^p<0.01 vs WT mice. Individual symbols in the bar graphs (F, H, J, L) represent individual mice. All data represent mean ± SEM. Statistical analysis was done by t-test.

**Supplemental Table S1.**
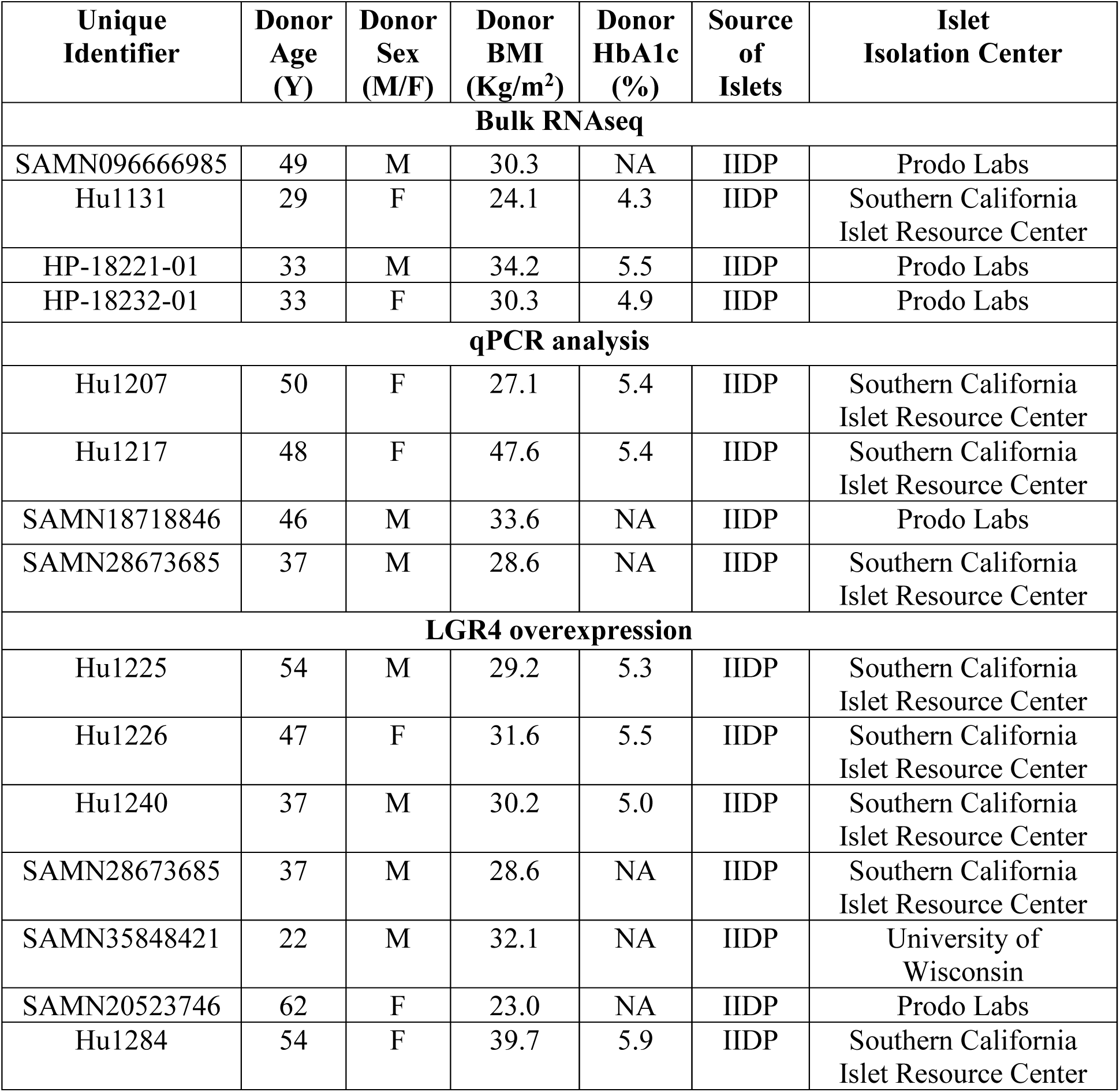
Donor characteristics associated with cadaveric human islets.

**Supplemental Table S2.** Reagents.

| Reagent | Company | Catalog # | Dilution |
| --- | --- | --- | --- |
| **Peptides** |  |  |  |
| Human-TNF-α | R&D Systems | 210-TA-020/CF | 13.16 ng/ml |
| Human-INF-γ | R&D Systems | 285-IF-100/CF | 50 ng/ml |
| Human-IL-1β | R&D Systems | 201-LB-005/CF | 0.72 ng/ml |
| Mouse-TNF- $\alpha$ | R&D Systems | 410-MT | 3.704 ng/ml |
| Mouse-INF-γ | R&D Systems | 485-MI | 118.62 ng/ml |
| Mouse-IL-1β | R&D Systems | 401-ML | 0.091 ng/ml |
| TRAF6 Inhibitor Peptide Set | Novus Biologicals | NBP2-26506 | 30 μM |
| **Antibodies** |  |  |  |
| p-PHH3-antibody | Millipore Sigma | 06-570 | 1:500 |
| BrdU | Abcam | 6326 | 1:500 |
| p-NFkB (Ser468) | Thermo Fisher Scientific | PA5-99380 | 1:200 |
| Cleaved Caspase-3 | Cell Signaling Technologies | 966-1S | 1:200 |
| Leucine Rich Repeat G Protein-Coupled Receptor 4 (LGR4) | R&D Systems | MAB7750 | 1:50 |
| Cre Recombinase | Cell Signaling Technologies | 15036 | 1:400 |
| Insulin | Abcam | ab96135356 | 1:800 or 1:1000 |
| Alexa Fluor 488 Goat α-Rabbit | Life Technologies | A11034 | 1:250 |
| Alexa Fluor 594 Goat α-Guinea Pig | Life Technologies | A11076 | 1:250 |
| Alexa Fluor 594 Goat α-Rabbit | Life Technologies | A11037 | 1:250 |
| Alexa Fluor 488 Goat α-Guinea Pig | Life Technologies | A11073 | 1:250 |
| Alexa Fluor 488 Goat α-Mouse | Life Technologies | A11029 | 1:250 |
| HRP Donkey α-Guinea Pig | Jackson ImmunoResearch | 706-035-148 | 1:500 |
| **Reagent Kits** |  |  |  |
| DeadEnd Fluorimetric TUNEL Kit | Promega | G3250 | NA |
| Mouse insulin ELISA kit | Mercodia | 10-1249-01 | NA |
| DAB Peroxidase Substrate Kit | Vector Labs | SK-4100 | NA |
| **Reagents** |  |  |  |
| DAPI (4',6-Diamidino-2-Phenylindole) | Life Technologies | D3571 | 1:200-2000 |
| SC siRNA negative control | Thermo Fisher Scientific | 4390843 | 25 or 125nM/L |
| *Lgr4* siRNA | Thermo Fisher Scientific | AM16704 | 25nM/L |
| *Rank* siRNA | Dharmacon Reagents | J103986170005 | 100nM/L |
| Lipofectamine RNAiMax | Thermo Fisher Scientific | 13778030 | 1:11 |
| iTaq Universal SYBR® Green Supermix | BIO-RAD | 1725124 | 1:1 |
| Taqman™ Fast Advanced Master Mix | Thermo Fisher Scientific | 4444556 | 1:1 |
| Ammonium pyrrolidinedithiocarbamate (PDTC) | Sigma-Millipore | P8765-1G | 250µM |
| Collagenase P | Roche | 11249002001 | NA |
| BrdU | Thermo Fisher Scientific | B23151 | 10µM |
| D-(+)-Glucose | Sigma-Millipore | G7021-1KG | 5.5mM, 11.5mM (*in vitro*), 2g/kg body weight (*in vivo*) |
| Humulin (Recombinant Human Insulin) | Eli Lilly | HI-213 | 1.5U/g |
| Streptozotocin (STZ) | Sigma-Millipore | 572201-250MG | 40mg/kg |
| Rodent Diet with 60 kcal% Fat | Research Diets, Inc. | D12492 | NA |
| Hematoxylin | Sigma-Aldrich | HHS16-500 | NA |
| Hydrogen Peroxide | Sigma-Aldrich | H1009 | 1:10 |
| Sodium chloride (NaCl 129mM) | Sigma-Aldrich | S7653 | NA |
| Potassium chloride (KCl 4.8 mM) | Sigma-Aldrich | P9333 | NA |
| Calcium chloride dihydrous (CaCl<sub>2</sub> 2H<sub>2</sub>O 2mM) | Sigma-Aldrich | C1016 | NA |
| Sodium bicarbonate (NaHCO<sub>3</sub> 24 mM) | Sigma-Aldrich | C5761 | NA |
| Potassium dihydrogen phosphate (KH<sub>2</sub>PO<sub>4</sub> 1.2mM) | Sigma-Aldrich | 10487 | NA |
| Magnesium sulfate heptahydrate (Mg<sub>2</sub>SO<sub>4</sub> 7H<sub>2</sub>O 1.2 mM) | Sigma-Aldrich | 1374361 | NA |
| HEPES (6 mM) | Corning | 25-060-CI | NA |
| BSA (0.2%) | Gemini | 700-100P | NA |
| Histopaque-1077 | Sigma-Aldrich | 10771-100ML | NA |
| Fluoro-Gel mounting<br>media | Electron<br>Microscopy<br>Sciences | 17985-10 | NA |

**Supplemental Table S3.**
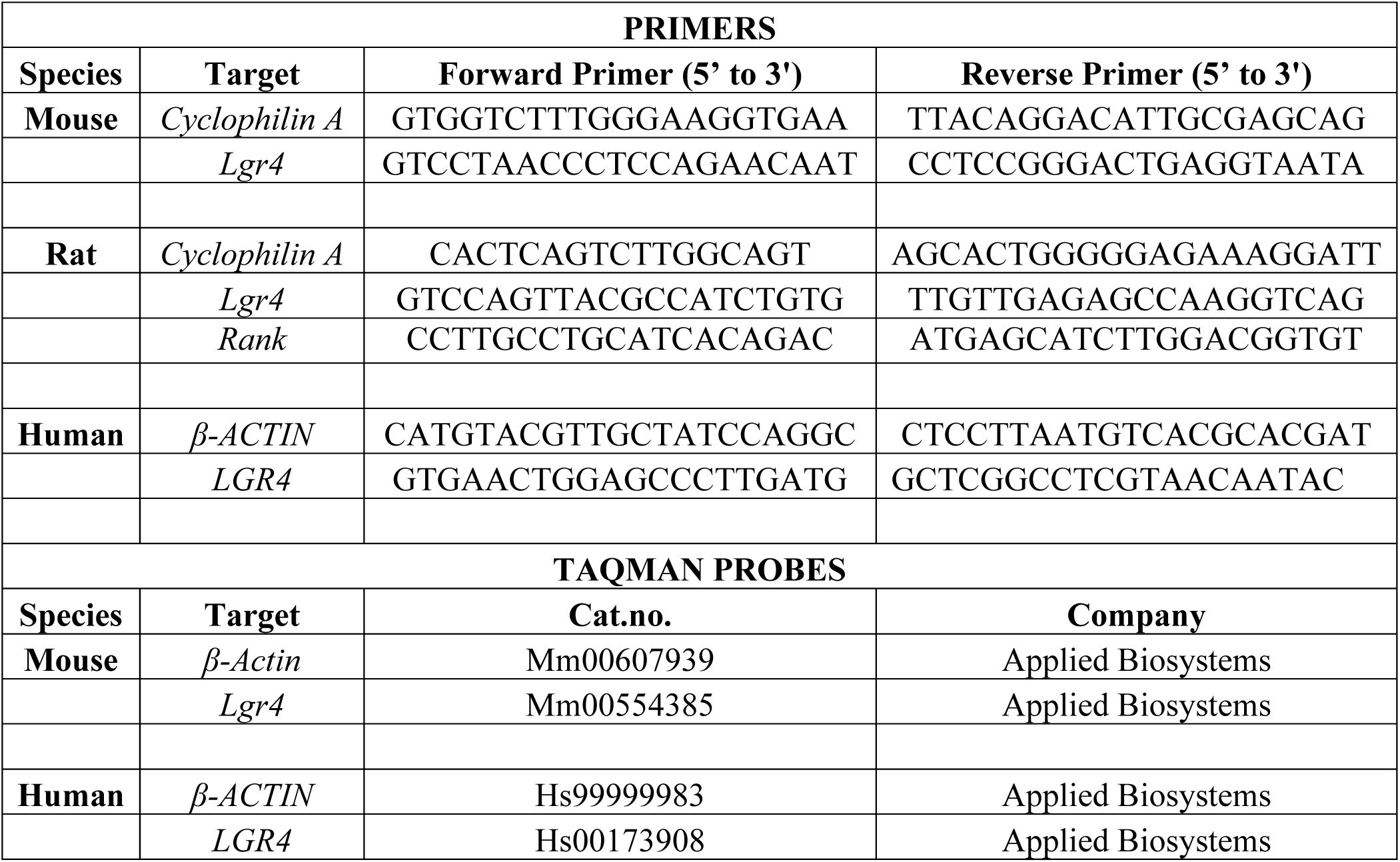
qPCR Primers and Taqman Probes.

